# *MAPK14*/p38α Shapes the Molecular Landscape of Endometrial Cancer and promotes Tumorigenic Characteristics

**DOI:** 10.1101/2024.06.25.600674

**Authors:** Sayali Joseph, Xingyuan Zhang, Gaith Droby, Di Wu, Victoria Bae-Jump, Scott Lyons, Angie Mordant, Allie Mills, Laura Herring, Blake Rushing, Jessica Bowser, Cyrus Vaziri

**Affiliations:** Department of Pathology and Laboratory Medicine, University of North Carolina, Chapel Hill, NC 27599, USA; Curriculum in Genetics and Molecular Biology, University of North Carolina, Chapel Hill, NC 27599, USA; Department of Biostatistics, University of North Carolina, Chapel Hill, NC 27599, USA; Lineberger Comprehensive Cancer Center, University of North Carolina, Chapel Hill, NC 27599, USA; Department of Pharmacology, UNC Proteomics Core Facility, University of North Carolina, Chapel Hill, NC 27599, USA; Nutrition Research Institute, University of North Carolina at Chapel Hill, Kannapolis, NC 28081, USA; Department of Nutrition, University of North Carolina at Chapel Hill, Chapel Hill, NC 27599, USA

## Abstract

The molecular underpinnings of <u>H</u>igh <u>G</u>rade <u>E</u>ndometrial <u>C</u>arcinoma (HGEC) metastatic growth and survival are poorly understood. Here we show that ascites-derived and primary tumor HGEC cell lines in 3D spheroid culture faithfully recapitulate key features of malignant peritoneal effusion and exhibit fundamentally distinct transcriptomic, proteomic and metabolomic landscapes when compared with conventional 2D monolayers. Using genetic screening platform we identify *MAPK14* (which encodes the protein kinase p38α) as a specific requirement for HGEC in spheroid culture. *MAPK14*/p38α has broad roles in programing the phosphoproteome, transcriptome and metabolome of HGEC spheroids, yet has negligible impact on monolayer cultures. *MAPK14* promotes tumorigenicity *in vivo* and is specifically required to sustain a sub-population of spheroid cells that is enriched in cancer stemness markers. Therefore, spheroid growth of HGEC activates unique biological programs, including p38α signaling, that cannot be captured using 2D culture models and are highly relevant to malignant disease pathology.

## Introduction

Endometrial Cancer (EC) is one of the few solid tumors with rising incidence and mortality rates. <u>H</u>igh <u>G</u>rade <u>E</u>ndometrial <u>C</u>arcinomas (HGECs), which include high grade endometrioid EC and serous carcinomas, are the most aggressive forms of EC, accounting for ∼20-30% of cases and 50% of all EC deaths. More than 90% of patients develop inherent and/or acquired chemoresistance and few receive durable responses.^1, 2, 3 4^ While HGEC share genetic features with many other cancers, tumorigenic characteristics of HGEC, including responses to therapy, are generally not informed by other tumor types harboring similar driver mutations. To develop better treatment strategies for HGEC, it is necessary to define specific molecular determinants of the disease. The use of preclinical models that faithfully recapitulate HGEC disease pathobiology will be critical to elucidating genes and pathways that drive HGEC.

One hallmark of gynecological cancers (some HGECs and ovarian cancer) is peritoneal dissemination, which involves vasculature-independent local invasion of pelvic and abdominal organs.^5^ Approximately one-third of EC patients develop malignant peritoneal effusion (ascites) and abdominal fluid containing clusters of metastatic cancer cells termed ‘spheroids’. The unusual anchorage-free and vascular-independent conditions in which spheroids arise may drive unique tumorigenic properties^6^ such as intratumor heterogeneity^7, 8^ chemoresistance, dormancy and recurrence.^9, 10, 11, 12, 13, 14^

To better model key features of peritoneal metastases, it is feasible to culture cancer cells as 3D spheroids.^15, 16, 17, 18^ Multicelllar spheroids grown *ex vivo* closely recapitulate the features of metastatic spheroids in ascites including cell-cell interactions^19^, differentiation,^20^ chemical gradients, a proliferation gradient (decreasing from periphery to center)^16, 17, 21^ and a central zone of dead cells.^22, 23, 24, 25^ For several malignancies, including ovarian cancer, therapeutic responses in 3D spheroids correlate better with *in vivo* responses when compared with 2D cultures.^26, 27^ Another feature of ovarian cancer 3D spheroids is that they are enriched in cancer stem cells (CSC). CSCs are multipotent cells that self-renew and confer tumorigenic features such as tumor seeding, metastatic spread and chemoresistance.^28, 29^ Identifying CSC is challenging because of their plasticity and low abundance.^29^ Cell surface markers (including CD133) are often associated with CSCs^30, 31, 32, 33, 34^ but not causally linked to stemness. CSC biology remains poorly understood and CSCs have not been molecularly defined in HGEC. Our current understanding of genes and pathways that sustain HGEC is primarily based on 2D cultures, not 3D model systems which reflect the pathobiology of human disease most closely. Due to the lack of studies with patient-relevant model systems, biology essential to HGEC may have gone unnoticed and may help explain why past studies have not readily translated into improved therapies for EC.

Here we sought to identify mechanisms that sustain spheroid growth and specify HGEC tumorigenic characteristics, with a focus on stress tolerance pathways. Neoplastic cells experience stress and genotoxicity from many sources including oncogenes, metabolites, aberrant mitotic programs, and DNA repair defects.^35^ We reasoned that cancer cells in spheroids experience unique stresses that do not exist in 2D cultures. For example, spheroids develop a central zone of dying cells which may be caused by hypoxia, glucose deficiency and accumulation of toxic waste products.^15, 16, 17, 36^ Because neoplastic cells arise in stressful environments they often rely on adaptive stress-tolerance mechanisms to survive.^37, 38^ Thus, stress-tolerance mechanisms are enabling features and molecular vulnerabilities of neoplastic cells and could provide opportunities for therapy.

Using comprehensive transcriptomic, proteomic and metabolomic profiling approaches, we show that HGEC spheroids exhibit fundamentally distinct biology from HGEC cells in 2D culture. Using an unbiased screening approach, we identify *MAPK14* as a key gene that is required for HGEC spheroid growth but is dispensable for monolayers. *MAPK14* encodes p38α, a non-essential protein kinase and mediator of stress responses, whose roles in cancer are controversial.^39^ We show that *MAPK14*/p38α plays a key role in establishing the molecular landscape of HGEC only in multicellular spheroids and promoting tumorigenic characteristics.

## Results

### HGEC spheroids are enriched for markers of cell stress and cancer stemness

We developed conditions for growing HEC-50B cells (derived from ascites of a recurrent stage 3 tumor^40^) as 3D spheroids. The resulting spheroids comprised up to 20,000 cells with 3D cell-cell contacts recapitulating interactions of neoplastic cells within metastatic tumors (Figure 1A). HEC-50B spheroids contained proliferating outer zones and necrotic centers (Figure 1A, 1B). When compared with 2D monolayer cultures, spheroids expressed higher levels of DNA replication stress and DNA damage markers including pRPA, γH2AX, pCHK1 and pCHK2 (Figure 1C) and had higher levels of hypoxia (Figure 1D) and reactive oxygen species (Figure 1E). Spheroids were also enriched for protein and mRNA markers of cancer stem cells (CSCs) including CD133, SOX2, NANOG, KLF4, and OCT4 (Figures 1F, 1G). Thus, spheroids have distinct properties from 2D cultures and represent a pathologically-relevant system for identifying genes that sustain HGEC.

**Figure 1.**
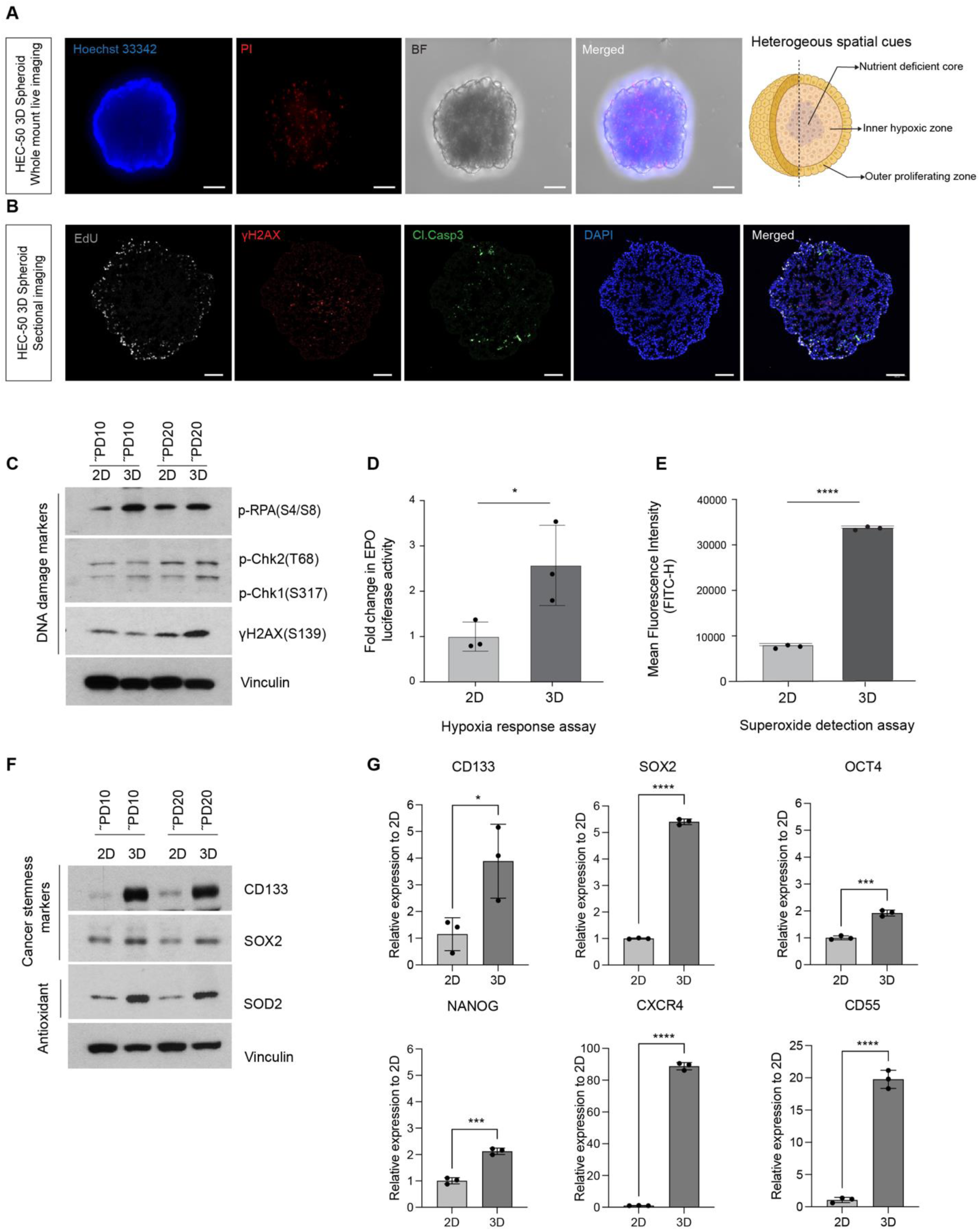
HEC-50B spheroids are enriched for markers of cell stress and cancer stemness. **(A)** Whole mount images of live HEC-50B spheroids stained with Hoechst 33342 and Propidium Iodide (PI). **(B)** Representative immunofluorescence staining images of HEC-50B spheroid cryosections showing EdU labeling, γH2AX and cleaved caspase 3 (Cl. Casp3). **(C)** Immunoblot showing relative levels of the indicated DDR markers after 10 and 20 population doublings in monolayer and spheroid culture. **(D)** Relative levels of EPO promoter-driven luciferase reporter gene activity in HEC-50B monolayers and spheroids. **(E)** Quantification of superoxide levels in monolayer and spheroid cultures of HEC-50B cells. **(F)** Immunoblot analysis of CSC markers and ROS scavenger (SOD2) in HEC-50B cells after 10 and 20 population doublings in monolayer and spheroid culture. **(G)** RT-PCR analysis showing relative levels of the indicated CSC markers in HEC-50B spheroids. In (A)–(B), HEC-50B spheroids were cultured for a duration of 8 days in ULA U-bottom-96-well plates. In (D), (E) and (F) unpaired two-tailed t-test was used to determine statistical significance. ∗p < 0.0332, ∗∗p < 0.0021, and ∗∗∗p < 0.0002, ∗∗∗∗p<0.0001. All data are shown as mean ± SD.

### CRISPR screening identifies MAPK14 as a dependency of HGEC cells in spheroid culture

We developed a genetic screening platform to identify putative genes that sustain HEC-50B spheroids but are dispensable for monolayer growth. We performed pooled CRISPR-CAS9 dropout screens using a cell stress and DNA damage response (DDR)-focused sgRNA library^41^ (Figure 2A).

**Figure 2.**
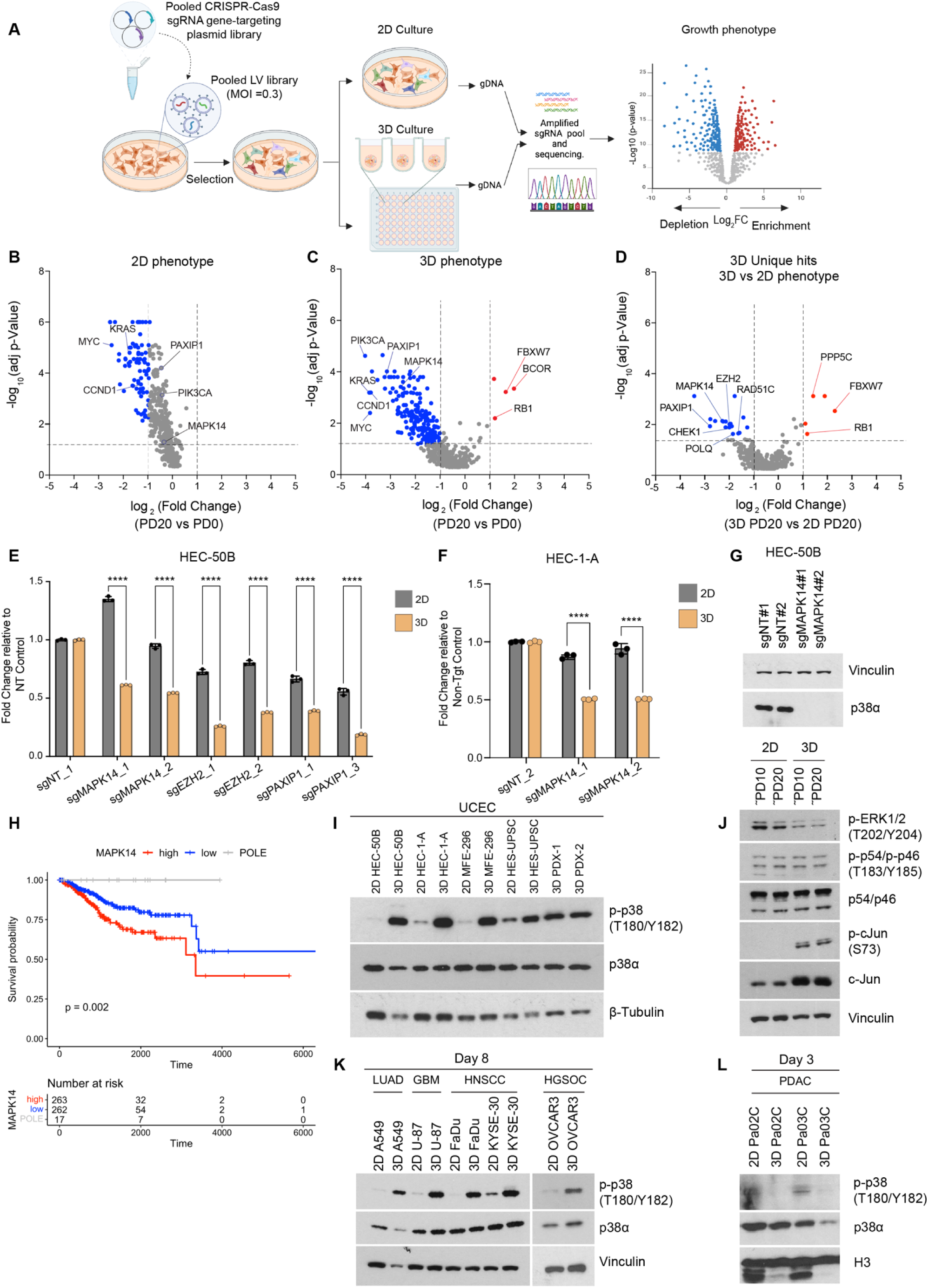
CRISPR screening identifies *MAPK14* as a dependency of HGEC in spheroid culture. **(A)** Schematic diagram of CRISPR screen workflow created using Birender.com. **(B)-(C)** Volcano plots depicting significant dropout (blue dots) and enrichment (red dots) of sgRNAs following 20 population doublings in monolayer (2D) and spheroid (3D) culture, relative to respective PD0. **(D)** Volcano plot showing sgRNAs that exhibited dropout or enrichment uniquely in spheroid (3D) cultures relative to monolayers (2D). **(E)** Effects of sgRNAs targeting *MAPK14*, *EZH2*, and *PAXIP1* (or non-targeting sgRNA, sgNT) on proliferation of HEC-50B spheroids after 14 days of culture. **(F)** Effects of *MAPK14*-directed sgRNA on proliferation of HEC-1-A cells in 3D spheroid culture. **(G)** Immunoblot confirming ablation of p38α expression by two independent sgRNAs targeting *MAPK14*. **(H)** Kaplan-Meier curves showing overall survival probability of endometrial cancer patients (TCGA uterine cancer cohort) expressing high (upper quartile) or low (lower quartile) levels of *MAPK14* mRNA. **(I)** Immunoblot showing levels of phospho-p38α (T180/Y182) in extracts from the indicated UCEC cell lines or PDX when grown in monolayer or spheroid culture. **(J)** Immunoblots showing effect of spheroid culture on levels of phosphorylated ERK1/2 (T202/Y204), JNK (T183/Y185) in HEC-50B cells. **(K)** Immunoblots showing levels of phospho-p38α (T180/Y182) in LUAD (A549), GBM (U87), HNSCC (FaDu, KYSE-30), HGOSC (OVCAR3), and (L) PDAC (Pa02C, Pa03C) cells when grown in monolayer (2D) or spheroid (3D) culture. In (B)–(D), Kolmogorov-Smirnov test was used to determine significant sgRNA enrichment or depletions. In (E) and (F) Ordinary Two-way Anova with Tukey’s test was used to determine statistical significance. In (H) Log-rank tests were used to infer statistical significance between high and low *MAPK14* groups. In plots (E-F) depict ± SD. ∗p < 0.0332, ∗∗p < 0.0021, and ∗∗∗p < 0.0002, ∗∗∗∗p<0.0001.

As expected, there was no significant depletion or enrichment of our internal control Non-Targeting (NT) sgRNAs over 20 population doublings (PDs), either in 2D or 3D cultures (Figure S1A). In contrast, 63 of the 504 sgRNAs in our sgRNA library were depleted in monolayers and spheroids (PD20 vs PD0), suggesting that their corresponding genes were essential for both 2D and 3D growth (Figure 2B, Figure 2C, Figure S1B). The 63 common essential genes included several protooncogenes (e.g. *KRAS* and *MYC*) and core components of DDR pathways (e.g. *ATM* and *RAD51C*). We identified 9 genes that were required for 2D growth but dispensable for 3D growth whereas 102 genes were necessary for growth as spheroids but not monolayers. These results suggest that cell growth in 3D creates a greater dependency on stress and DDR responses as compared with 2D monolayer culture (Figure S1B-C).

After 20 PDs, sgRNAs targeting *FBXW7* and *RB1* (known tumor-suppressor genes for endometrial cancer) were enriched in 3D-spheroids but not in monolayers (Figure 2D) suggesting that *FBXW7* and *RB1*-deficiencies promote growth of HEC-50B cells specifically when in spheroid culture. Conversely, *PIK3CA*, an important driver oncogene in endometrial cancer was essential for spheroid growth but dispensable for monolayers. Therefore, HGEC cells in 2D and 3D culture have fundamentally different requirements for tumor suppressor genes, oncogenes and DDR factors.

To identify genetic dependencies unique to 3D growth, we compared dropout of sgRNAs between spheroid and monolayer cultures at PD20 (Figure 2D). Those analyses identified 18 genes (*PIK3CA*, *PAXIP1*, *SMAD4*, *CCND1*, *CDK2*, *MAPK14*, *CDC5L*, *ID2*, *CHEK1*, *EZH2*, *RINT1*, *KDM2A*, *POLQ*, *SMAD5*, *LIG3*, *RAD51C*, *SMAD1*, *CHD8*) as requirements for 3D growth. We individually validated some of these spheroid-specific dependencies. As shown in Figure 2E, sgRNAs targeting *MAPK14*, *PAXIP1* or *EZH2* preferentially inhibited growth of HEC-50B cells in spheroid cultures when compared with 2D monolayers. Similar results were seen when we ablated *MAPK14* in HEC-1-A cells (Figures 2F, 2G, S1D). Interestingly, high expression of *MAPK14* and *EZH2* (but not of *PTIP*) correlates with low survival probability of HGEC patients (Figures 2H, S1F, S1G), potentially consistent with pro-tumorigenic roles. Cell cycle profiles of HEC-50B cells derived from WT and *MAPK14^-/-^* spheroids were indistinguishable (Figures S1H, S1I). Moreover, *MAPK14*-loss was associated with decreased necrosis and apoptosis (Figures S1J, S1K). We conclude that the effect of *MAPK14*-loss on spheroid growth is not due to overt cell cycle defects or loss of cell viability.

*MAPK14* encodes the p38α Stress-Activated Protein Kinase, a member of the Mitogen-Activated Protein Kinase (MAPK) superfamily. We observed higher levels of activation-associated p38α phosphorylation^39, 42^ in 3D spheroids when compared with 2D monolayer cultures in several EC cell lines including HEC-1-A, MFE-296, and HES-UPSC (Figure 2I). We also detected high-level activation of p38α kinase in 3D cultures of EC patient-derived xenografts (PDX) (Figure 2I). The p38-related protein kinases ERK and JNK were not preferentially phosphorylated in HGEC spheroid cultures when compared with monolayers (Figure 2J). We also observed 3D-specific activation of p38α in cell lines derived from High Grade Ovarian Serous Carcinoma (HGOSC), Lung Adenocarcinoma (LUAD), Glioblastoma (GBM), and Head and Neck Squamous Cell Carcinoma (HNSCC) (Figure 2K). However, in pancreatic ductal adenocarcinoma (PDAC, a solid tumor that shares many HGEC oncogenes and tumor suppressors) cells, p38α phosphorylation was reduced in 3D cultures when compared with 2D monolayers (Figure 2L). Therefore, p38α phosphorylation is a feature of many cancer cells growing in 3D culture.

### MAPK14 establishes transcriptional programs involved in chemokine signaling and stress responses in HEC-50B spheroids

To identify candidate p38α-mediated gene expression programs involved in spheroid growth we compared transcriptional profiles of *MAPK14^+/+^* and *MAPK14^-/-^* HEC-50B cells in 2D and 3D culture. The Principal Component Analysis (PCA) plot in Figure 3A shows that 3D culture drove the largest global changes in the transcriptome in both *MAPK14^+/+^* and *MAPK14^-/-^* HEC-50B cells. *MAPK14* status had a negligible effect on the transcriptional profiles of monolayers, yet caused significant changes in the spheroid transcriptome. First we defined the differences between transcriptomes of wild-type (*MAPK14^+/+^*) cells in monolayer and spheroid culture. As shown in Figure 3B, 2422 genes were upregulated and 1484 genes were downregulated in 3D cultures when compared with monolayers. Next, we determined which of the transcriptional changes induced by spheroid growth were *MAPK14*-dependent. Figure 3C shows that 504 transcripts were differentially expressed between *MAPK14^+/+^* and *MAPK14^-/-^* spheroids, with 275 mRNAs repressed and 229 induced in *MAPK14^-/-^* spheroids when compared with wild-type controls. Of the genes that were differentially expressed between *MAPK14^+/+^* spheroids and *MAPK14^+/+^* monolayers, 256 (8.27%) were de-regulated in *MAPK14*^-/-^ spheroids. Therefore, ∼8.3% of the endometrial cancer cell transcriptional program induced by 3D spheroid culture is dependent on *MAPK14* (Figure 3E-F). In contrast with cells in spheroid culture, *MAPK14*-deficiency only altered expression of 280 mRNAs in 2D cultures (223 mRNAs were repressed and 57 were induced in *MAPK14^-/-^* monolayers when compared with wild-type – Figure 3D). Figure S2A provides an aerial view of all the mRNAs that were differentially expressed between 2D and 3D cultures, and indicates how their expression levels were impacted by *MAPK14*-deficiency.

**Figure 3.**
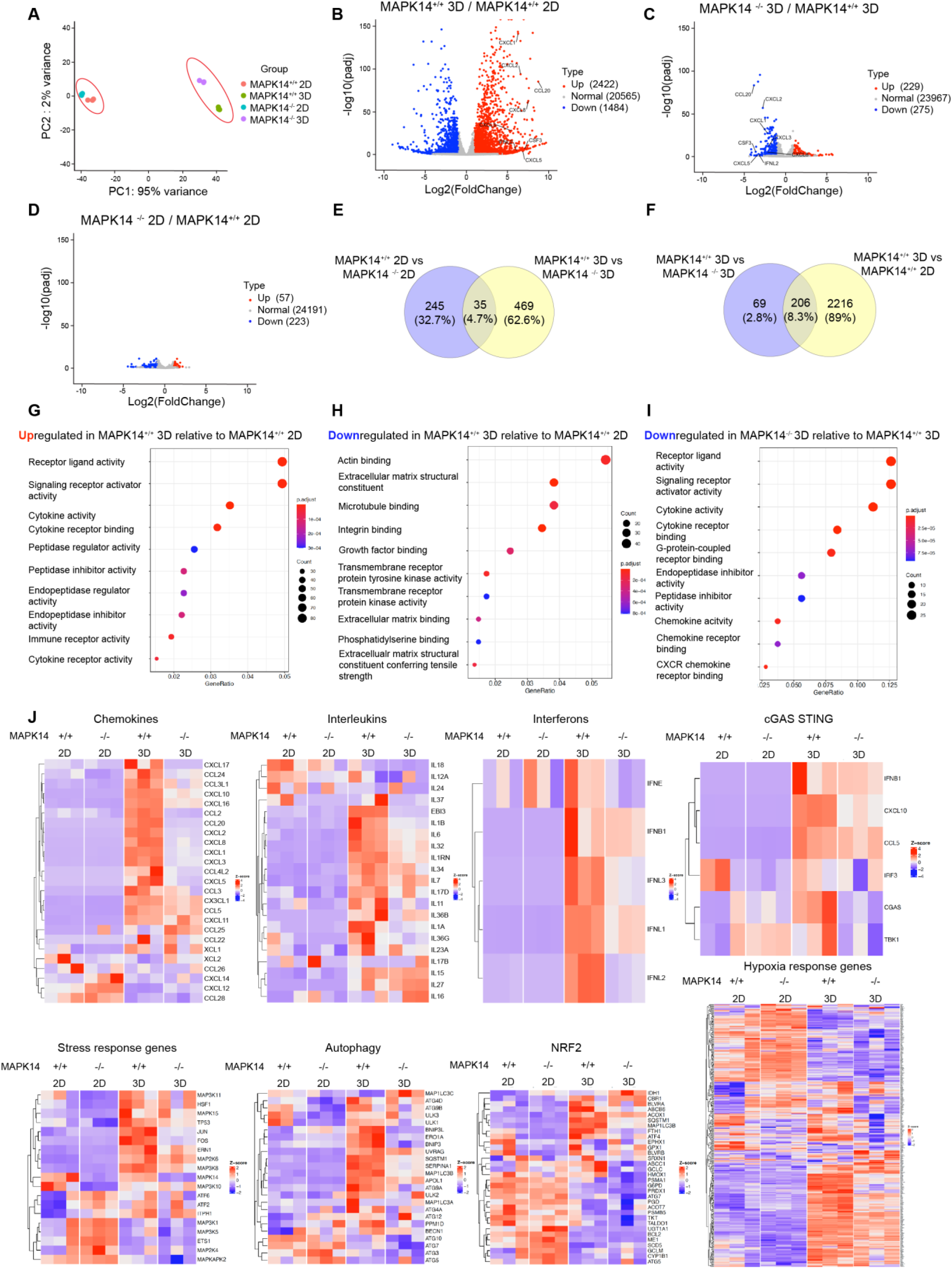
M*A*PK14 establishes transcriptional programs involved in chemokine signaling and stress responses in HEC-50B spheroids. **(A)** Principal component analysis (PCA) of bulk RNA-Seq data from *MAPK14^+/+^* and *MAPK14^-/-^* HEC-50B cells cultured as 2D monolayers and 3D spheroids. **(B-D)** Volcano plots showing log2fold changes (FC) in levels of individual expressed genes (red dots: increases; blue dots: decreases) against the −log10 adjusted p-value for the indicated pairwise comparisons of experimental samples. **(E-F)** Venn diagrams showing the numbers of differentially expressed genes (DEGs) in the indicated comparison groups, and the percentage of the transcriptomes that either overlap or have exclusive gene expression. **(G-I)** GO enrichment analyses indicating molecular functions of gene sets significantly enriched in DEGs of the indicated comparisons sets. **(J)** Heatmaps showing the effect of *MAPK14* genotype and spheroid culture on expression levels of individual mRNAs within the indicated pathways and processes

Our gene set enrichment analysis showed that chemokines, pro-tumorigenic cytokines, Interleukins, Interferon-induced genes, and endopeptidase inhibitors were the top gene categories that were up-regulated in wild-type spheroids when compared with monolayers (Figures 3G and S2B). Gene sets involved in cell adhesion factor binding, extracellular matrix organization and RhoGTPase signaling were downregulated in *MAPK14^+/+^* spheroids relative to *MAPK14^+/+^* monolayers (Figures 3H, S2C). Many of the genes expressed during 3D growth were dependent on *MAPK14* (Figures 3H, 3I, S2D). Figures 3J and S2E show the impact of 3D culture and *MAPK14* on the relative expression patterns of specific genes associated with some of these signaling pathways including chemokines, cytokines, IFNs, autophagy, NRF2, and responses to hypoxia. We conclude that p38α is broadly required for activation of transcriptional programs associated with 3D growth.

### The phosphoproteome of HEC-50B cells in spheroid culture is p38α-dependent

To further elucidate roles of p38α signaling in HGEC spheroid biology, we profiled the proteomes and phosphoproteomes of WT and *MAPK14^-/-^* cells in monolayer and spheroid culture. PCA revealed that spheroid culture elicited large changes in the proteome and the phosphoproteome when compared with monolayers (Figures 4A, 4B). *MAPK14*-deficiency had relatively little effect on the proteome or phophoproteome in monolayers. However, proteome and phosphoproteome profiles were more divergent when comparing *MAPK14^+/+^* and *MAPK14^-/-^* cells in spheroid culture (Figures 4C, 4D, 4E). We compared the fold changes in protein levels (identified by global proteome profiling) with the mRNA fold changes (identified by bulk RNASeq) and confirmed that there was concordance between the results of our proteomic and transcriptomic experiments (Figures S3A, S3B, S3C).

**Figure 4.**
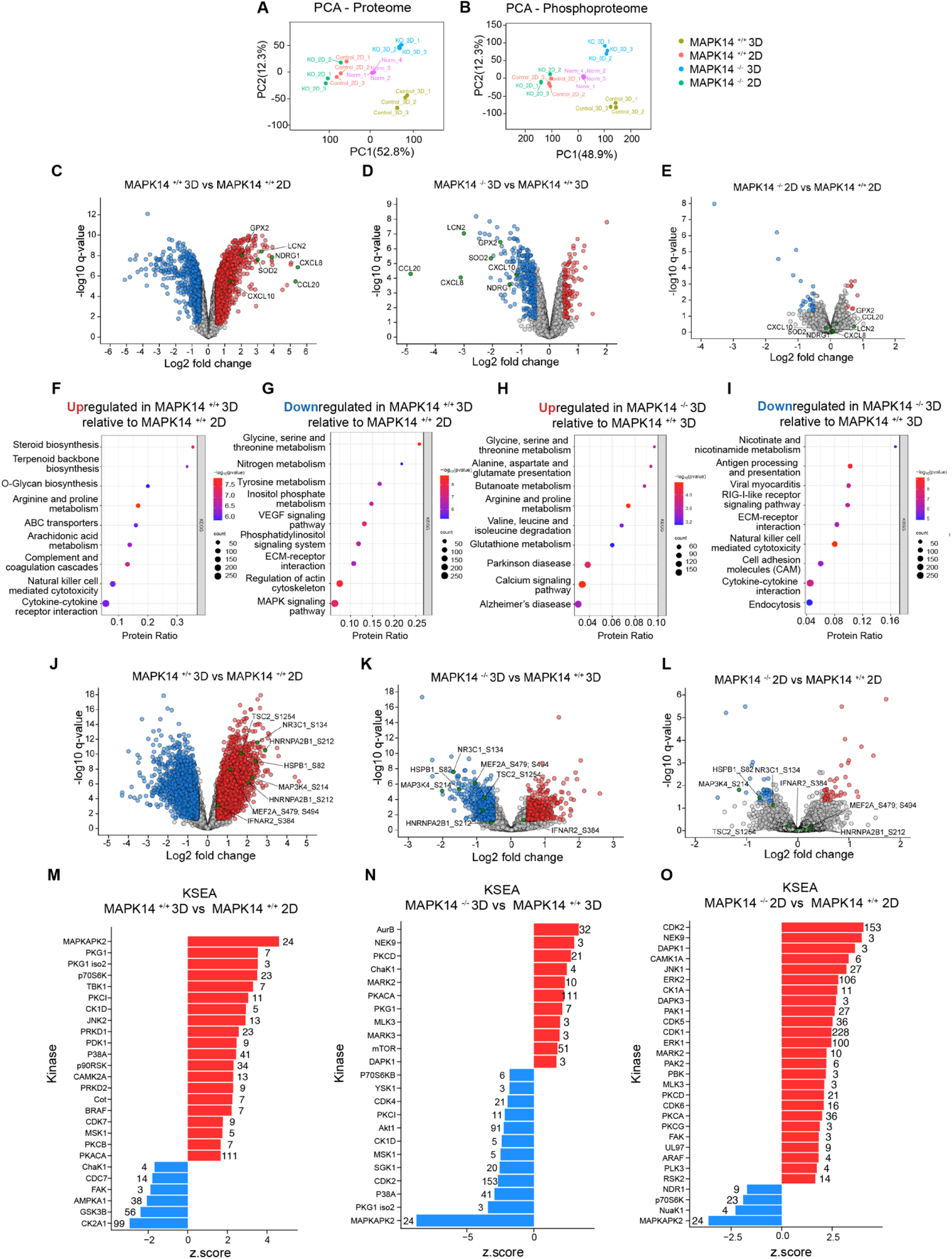
The phosphoproteome of HEC-50B cells in spheroid culture is p38α-dependent. **(A-B)** PCA plots of proteome (left panel) and phosphoproteome (right panel) data showing grouping of experimental samples from *MAPK14^+/+^* and *MAPK14^-/-^* HEC-50B cells cultured as monolayers and spheroids. **(C-E)** Volcano plots showing the log2 FC against the −log10 adjusted p value (the q-value) for differentially expressed proteins between the indicated sample pairs. Proteins with log2FC<-0.5 are indicated as blue dots, while proteins with log2FC>0.5 are indicated with red dots. A q value<0.05 was considered significant. **(F-I)** Bubble plots depicting KEGG pathway enrichment analyses of differentially expressed proteins either upregulated or downregulated, as indicated in the comparisons sets. **(J-L)** Volcano plots showing the log2 FC against the −log10 adjusted p value (q-value) for differentially expressed phosphopeptides between the indicated sample pairs. Phosphopeptides colored blue have a log2FC < −0.5 and a q-value < 0.05 while those colored red have a log2FC > 0.5 and a q-value < 0.05, which are considered biologically significant. **(M-O)** Results of KSEA showing all significantly-altered kinases with either low or high z-scores. Red bars indicate kinases with significantly increased activity while blue bars indicate kinases with significantly decreased activity. The numeral next to each bar specifies the number of substrates identified for the indicated kinase.

We observed an increase in levels of cytokine and chemokine signaling proteins in *MAPK14^+/+^* cells growing as spheroids when compared with monolayer culture (Figure S3D). IPA analysis showed that pathways related to cytokine signaling, cholesterol biosynthesis, NF-kB, NRF2-mediated oxidative stress response, and glutathione redox reactions were increased in spheroids compared with monolayers (Figure S3E). The spheroid-specific expression of chemokines such as IL8 and CCL20 was *MAPK14*-dependent at the protein level (Figure 4C).

KEGG pathway analysis also showed that proteins involved in metabolic processes such as steroid biosynthesis, arginine, and proline metabolism were upregulated in *MAPK14^+/+^* spheroids when compared with *MAPK14^+/+^* monolayers (Figure 4F) whereas proteins involved in glycine, serine, and threonine metabolism were downregulated during spheroid growth (Figure 4G). The downregulation of factors involved in glycine, serine and threonine metabolism in HGEC spheroids was *MAPK14*-dependent, suggesting that p38α promotes metabolic adaptation during 3D growth (Figure4H). KEGG pathway analysis confirmed that chemokine signaling and cell adhesion proteins were expressed in a *MAPK14*-dependent manner in spheroids (Figure 4I).

In *MAPK14^+/+^* cells, spheroid growth also led to robust reprogramming of the phosphoproteome. We detected upregulation of 3,886 phosphopeptides in *MAPK14^+/+^* spheroids when compared with *MAPK14^+/+^* monolayers (Figure 4J). We also determined the contribution of *MAPK14* to the phosphoproteome. In *MAPK14^-/-^* spheroids, we observed downregulation of 681 phosphopeptides and upregulation of 593 phosphopeptides when compared with *MAPK14^+/+^* spheroids (Figure 4K). In contrast, in *MAPK14^-/-^* monolayers, we only detected 36 phosphopeptides whose abundance decreased and 44 phosphopeptides whose abundance increased when compared with *MAPK14^+/+^* monolayers (Figure 4L). Therefore, *MAPK14*-deficiency broadly impacted the phosphoproteome of spheroids, yet was relatively inconsequential in monolayer cultures. Gene set enrichment analysis of differentially abundant phosphopeptides showed that regulation of RhoGTPase family signaling was *MAPK14*-dependent in spheroids (Figure S3F).

From Kinase Substrate Enrichment Analysis (KSEA), the top 5 protein kinase pathways preferentially activated in spheroids (when compared with monolayers) were MAPKAPK2 (MK2), PKG1, PKG1 isoform 2, S6K1, and TBK1 (Figure 4M). Notably, MK2 is a critical downstream target of p38α^42^. We also detected increased phosphorylation of MAP3K4 and MAP2K4 (known upstream activators in the p38α MAPK signaling module) in *MAPK14^+/+^* spheroids when compared with *MAPK14^+/+^* monolayers. KSEA identified the p38α target MK2 as the most downregulated protein kinase pathway in *MAPK14^-/-^* 3D cultures when compared with *MAPK14^+/+^* spheroids (Figure 4N). We also detected increased activation of AURB, NEK9, PKCD, ChaK1, and MARK2 kinase pathways in *MAPK14^-/-^* spheroids when compared with *MAPK14^+/+^* spheroids possibly reflecting a compensatory response to p38α-deficiency.

The abundance of phosphopeptides corresponding to known substrates of p38α (including transcription factors STAT1, STAT3 and NR3C1; protein kinases MAPKAPK5 and GSK3β) and MK2 (including molecular chaperones HSBP1, TSC2, and DNAJB1; transcriptional regulators and RNA processing factors RMB7, ZFP36L, HNRNPN0 and NELFE) was increased in 3D cultures when compared with monolayers (Figure S4A). Remarkably, the largest increases and decreases in phosphoprotein abundance caused by spheroid culture were all *MAPK14*-dependent, even when we did not restrict our analysis to known targets of p38α/MK2 signaling. (Figure S4B). Therefore, *MAPK14* plays a key role in specifying the global kinome of HGEC cells in spheroid culture.

We independently validated some of the key changes revealed by our transcriptional profiling and phosphoproteome analyses (Figure S5). We detected increased phosphorylation of p38α downstream targets such as MK2, GSK3β (T390),^43^ and NR3C1(S211)^44^ in spheroids when compared with monolayers. Downstream targets of MK2 such as Heat Shock Protein 27 (HSP27), at S82^45^ were also more highly phosphorylated in spheroids (Figures S5A, S5B). Consistent with our KSEA, we detected increases in the activation-associated phosphorylation of TBK1 (at S172)^46^ and S6K1 (at T389)^47^ in HGEC spheroids when compared with monolayers (Figures S5C, S5D). Total PKG1 levels were higher in spheroids (Figure S5C). Our metabolomic profiling (described below and in Figure 5) revealed that cGMP levels were 3.2-fold higher in *MAPK14^+/+^* spheroids than in *MAPK14^+/+^* monolayers, thereby reproducing the patterns of PKG1 signaling identified by our KSEA (Figure 4). The increases in phosphorylation and/or total protein levels of PKG1, TBK1 and S6K1 in spheroids when compared with monolayer cultures were *MAPK14*-independent (Figures S5C, S5D).

**Figure 5.**
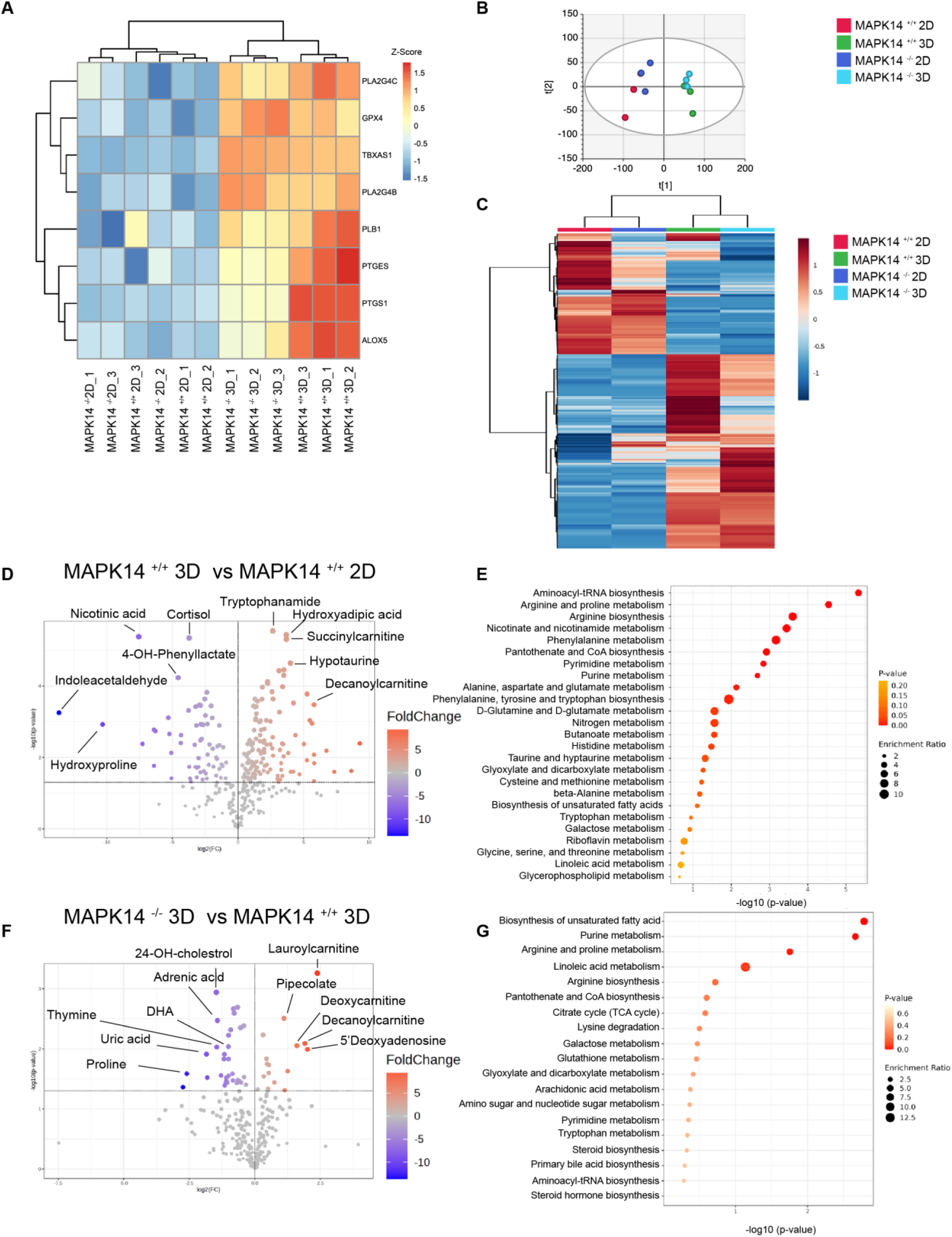
*MAPK14* specifies metabolic programs in HEC-50B spheroids. **(A)** Heatmap showing levels of proteins involved in arachidonic acid metabolism in HEC-50B monolayers and spheroids. **(B)** PCA plot of *MAPK14^+/+^* and *MAPK14^-/-^* HEC-50B cells cultured as spheroids and monolayers using all metabolomics peaks. **(C)** Heatmap of all in-house matched metabolites across all four groups. **(D)** Volcano plot of log2-adjusted fold change (FC) and −log(p-value) of in-house matched metabolites between *MAPK14^+/+^* monolayers and *MAPK14^+/+^* spheroids. A positive FC indicates metabolites increased in spheroids. **(E)** Pathway enrichment plot of differentially abundant metabolites with p<0.05 between *MAPK14^+/+^* monolayers and *MAPK14^+/+^* spheroids. **(F)** Volcano plot of log2-adjusted fold change (FC) and −log(p-value) of in-house matched metabolites between *MAPK14^+/+^* spheroids and *MAPK14^-/-^* spheroids. A positive FC indicates metabolites increased in *MAPK14^-/-^* samples. **(G)** Pathway enrichment plot of differentially abundant metabolites with p<0.05 between *MAPK14^+/+^* and *MAPK1^-/-^* spheroids.

Using TRRUSTv2 analysis^48^, we identified RELA, NFKB1, and STAT1 as candidate transcriptional activators of highly-expressed *MAPK14*-dependent genes in spheroids (Figure S5E). NF-кB and STAT1 are pro-inflammatory factors that could contribute to the expression of cytokines, chemokines, and interferons that we detected in spheroids (Figures 3, 4). Using immunoblotting with appropriate markers, we validated *MAPK14*-dependent activation of RELA and NFKB1 pathways in spheroids (Figures S5D, S5F). IKKβ and IKB phosphorylations associated with NF-кB pathway activation were induced in a *MAPK14*-dependent manner (Figures S5F, S5H), indicating that NFkB pathway activation in spheroids is p38α-mediated. Because there is extensive crosstalk between NF-kB, TBK1, STING and IFN signaling we also examined STING phosphorylation. Similar to IKKβ, STING phosphorylation (at Ser 366) was also induced in a *MAPK14*-dependent manner during spheroid growth (Figure S5D).

For STAT1 (a transcription factor which mediates responses to IFNs) both overall protein levels and phosphorylation at activating residues T701 and S727 were induced in spheroids, but independently of *MAPK14* (Figure S5D). The stress-responsive phosphoprotein NDRG1^49, 50, 51, 52^ and anti-oxidant factors Superoxide Dismutase 2 (SOD2)^53^ and Glutathione S-transferase alpha 1 (GSTA1)^54^ were also induced during spheroid culture in a *MAPK14*-dependent manner (Figure S5G). The transcription factor NRF2, which promotes expression of anti-oxidant and detoxification enzymes, and is involved in spheroid growth of lung cancer cells^55^ was more abundant in HGEC spheroids when compared with 2D cultures. The NRF2 target p62/SQSTM1 (which limits NRF2 degradation^56^) was also more highly phosphorylated in spheroids when compared with 2D monolayers (Figure S5C). However, the NRF2-KEAP pathway markers we examined were unaffected by *MAPK14* status (Figure S5C). Our results suggest that targets of p38α signaling (such as NF-kB1 and RELA)^39, 42^ specify the transcriptome of endometrial cancer spheroids and contribute to a proinflammatory environment.

### MAPK14 specifies metabolic programs in HEC-50B spheroids

From our proteome analysis (Figure 4F), some of the factors that were induced during spheroid culture regulate metabolism directly (e.g. SOD2, GSTA1; see Figure S5G) or indirectly (e.g. NRF2; Figure S3E). Consistent with the high inflammatory state revealed by the spheroid-specific transcriptome and proteome (Figures 3, 4), we observed elevated levels of proteins involved in arachidonic acid metabolism in spheroids (Figure 5A). To test the contributions of 3D growth and *MAPK14* status to the HGEC metabolome we measured levels of 7,923 metabolites extracted from *MAPK14^+/+^* and *MAPK14^-/-^* HEC-50B cells growing as monolayers and spheroids. PCA showed that the four experimental groups had distinct metabolite profiles (Figure 5B).

Figure 5C compares the relative abundances of 326 metabolites (which provide coverage of all major metabolic pathways) between *MAPK14^+/+^* and *MAPK14^-/-^* cells in both monolayer and spheroid culture. Of the 326 metabolites analyzed, 63 were downregulated and 131 were upregulated in *MAPK14^+/+^* spheroids when compared with *MAPK14^+/+^* monolayers (Figure 5C). Amino acid derivatives and acylcarnitines were among the metabolite classes most widely altered in the *MAPK14^+/+^* 3D versus *MAPK14^+/+^* 2D cultures (Figure 5D).

Spheroids showed an overall elevation of long chain acylcarnitines and decreases in short chain acylcarnitines, indicating mitochondrial dysfunction and uncoupling of fatty acid oxidation from oxidative phosphorylation.^57^ Pathway analysis of differentially-abundant metabolites (p<0.05) revealed that multiple metabolic pathways of amino acids as well as purine and pyrimidine metabolism were also significantly altered between monolayers and spheroid cultures (Figure 5E).

Comparison of *MAPK14^+/+^* and *MAPK14^-/-^* spheroids revealed that 16 metabolites were upregulated, and 33 metabolites were downregulated as a result of *MAPK14*-deficiency (Figure 5F). Acylcarnitines, nucleotides, and polyunsaturated fatty acids were among the most significantly altered metabolites in *MAPK14*-ablated spheroids (Figure 5G). Pathway analysis revealed that unsaturated fatty acid metabolism, arginine metabolism, glutathione metabolism, and purine/pyrimidine metabolism were the pathways that most significantly (p<0.05) differentiated *MAPK14^+/+^* and *MAPK14^-/-^* spheroids (Figure 5G). Our metabolomics and proteomics data both suggested a decrease in glycolysis and an increase in TCA cycle activity in *MAPK14^-/-^* spheroids when compared with *MAPK14^+/+^* spheroids (Figures S6A, S6B). We also detected altered metabolism of unsaturated fatty acids in spheroids lacking *MAPK14*. Notably, *MAPK14*-dependent metabolism of the eicosanoid precursor arachidonic acid (Figure S6C) may also contribute to a proinflammatory environment in spheroids.

We observed a significant decrease in cortisol in wild type 2D monolayers vs 3D spheroids as well as *MAPK14^-/-^* monolayers when compared with *MAPK14^-/-^* spheroids. Finally, we observed significant changes in several amino acid metabolites in *MAPK14^-/-^* monolayers when compared with *MAPK14^-/-^* spheroids. Amino acids in the methionine cycle (S-adenosylmethionine, S-adenosylhomocysteine, and homocysteine) were decreased while serine was increased, which is consistent with the rewiring of serine, glycine, and threonine metabolism indicated by the proteomics data. Taken together, the results of Figures 5 and S6 show that *MAPK14* contributes to the metabolic reprogramming of HGEC cells growing as multicellular spheroids.

### MAPK14 sustains sub-populations of cancer stem-like cells in spheroids

We used single-cell RNA (scRNASeq) to analyze the impact *MAPK14* loss on different spheroid sub-populations. We identified 11 transcriptionally-distinct cell types in HGEC spheroids (Figure 6A-B). Of those 11 sub-populations, ‘Cluster 3’ was the most affected by *MAPK14*-loss. Cells with a Cluster 3 signature were 14.5-fold less abundant in the *MAPK14^-/-^* spheroids when compared with *MAPK14^+/+^* spheroids (Figure 6C). A cell population represented by ‘Cluster 1’ was also less abundant (by 3.5-fold) in the *MAPK14^-/-^* samples relative to *MAPK14^+/+^* spheroids (Figure 6C). Cell populations represented by clusters 0 and 4, were more abundant (by 5.4-fold and 4.6-fold respectively) in the *MAPK14^-/-^* spheroids when compared with *MAPK14^+/+^* controls (Figure 6B). The abundance of sub-populations of cells represented by Cluster 2 and Clusters 5-9 was relatively unaffected by *MAPK14* loss.

**Figure 6.**
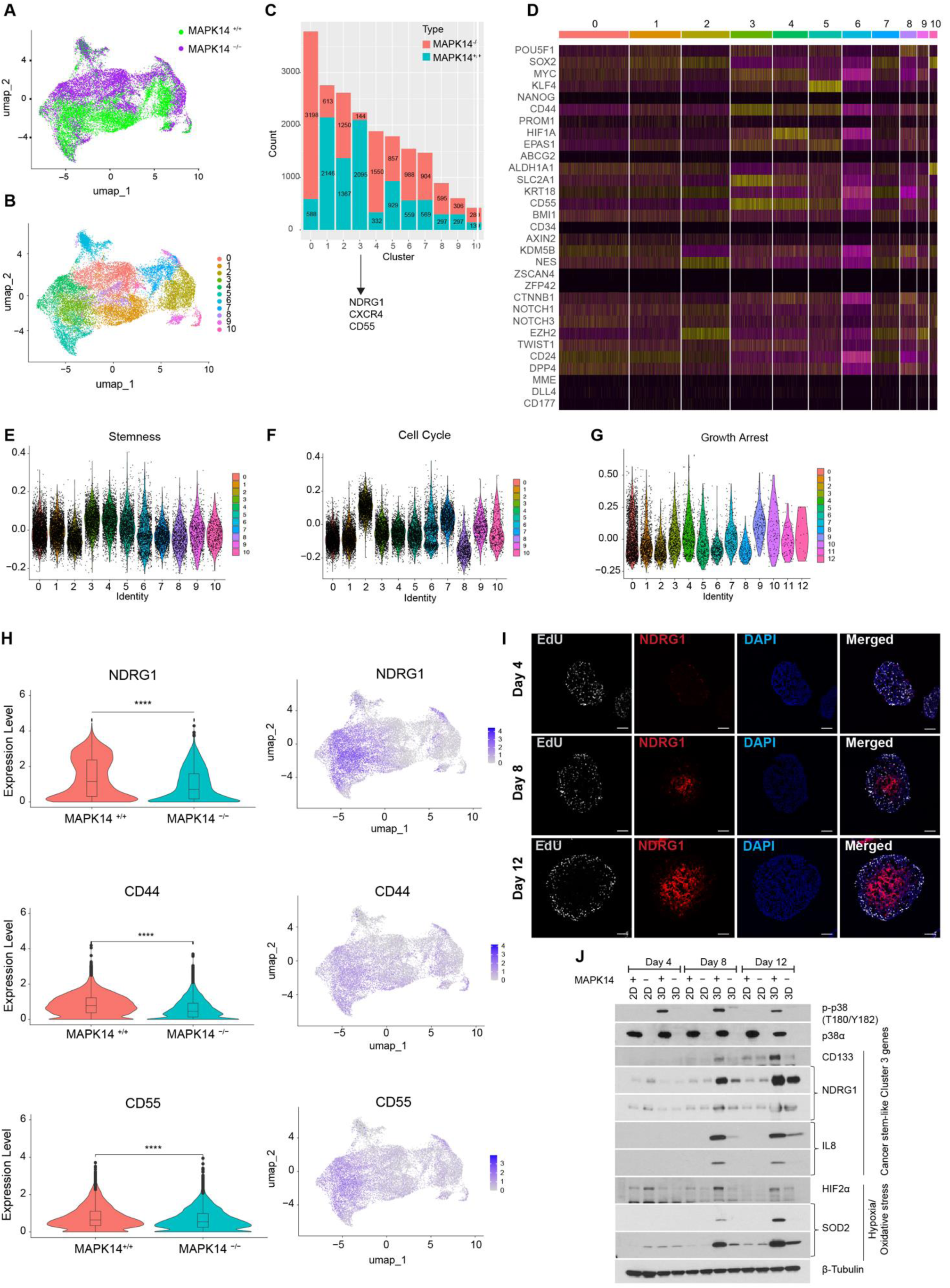
*MAPK14* sustains sub-populations of cancer stem-like cells in spheroids. **(A)** UMAP plot of single cell RNA-Seq data from HEC-50B spheroids colored according to *MAPK14^+/+^* and *MAPK14^-/-^* genotypes. **(B)** UMAP plot of scRNA-Seq data from *MAPK14^+/+^* and *MAPK14^-/-^* HEC-50B spheroids with colors indicating 11 transcriptionally-distinct clusters (identified by Seurat). **(C)** Stacked bar chart showing impact of *MAPK14* status on numbers of cells in each of the 11 transcriptionally-distinct clusters. **(D)** Heatmap showing expression levels of individual CSC markers, associated with endometrial cancers, in each of the 11 transcriptionally-distinct clusters **(E-G)** Plots showing relative levels of gene expression signatures for cancer stemness (E), DNA replication and cell cycle (F) and quiescence/growth arrest (G) in each of the 11 clusters identified by scRNA-Seq. **(H)** Violin plots showing relative expression levels of cancer-stemness associated markers *NDRG1*, *CD44*, and *CD55* mRNAs in *MAPK14^+/+^* and *MAPK14^-/-^* spheroids (left) and UMAP projections colored to indicate the clusters associated with expression of *NDRG1*, *CD44*, and *CD55* (right). **(I)** Immunofluorescence microscopy of HEC-50B spheroid sections showing distribution of cells expressing NDRG1 at days 4, 8, and 12 after seeding. Scale bars represent 100 μm. **(J)** Immunoblot analysis showing temporal expression patterns of the indicated proteins in *MAPK14^+/+^* and *MAPK14^-/-^* HEC-50B cells maintained in 2D or 3D culture for 4, 8, and 12 days. In the left panels of (H) Wilcoxon test was used to determine statistical significance. ∗∗∗∗p value <0.0001

To establish the biological features of cells represented by the 11 transcriptionally-distinct clusters we performed GO enrichment analysis of their individual transcriptomes. We were unable to reliably assign enrichment of any particular gene ontology groups to the Cluster 0 transcriptome. However, Cluster 4 cells were modestly enriched in genes related to ROS (Figure S7). Cells represented by Cluster 2 were characterized by high expression of DNA replication and cell cycle genes and therefore represent an actively-proliferating subset (Figure S1H). Instead, *MAPK14*-loss compromises maintenance of specific sub-populations in a multicellular spheroid.

The cells represented by Clusters 1 and 3 (the Clusters most severely depleted as a result of *MAPK14*-deficiency) were enriched in genes involved in epithelial maintenance and proteolytic processes (Cluster 1) and responses to hypoxia (Cluster 3) (Figure S7). The Cluster 3 transcriptome was also highly enriched for a cancer stemness gene expression signature^58^ (Figures 6D, 6E). CSCs exist in a quiescent growth-arrested state. The Cluster 3 transcriptome expressed low levels of cell cycle genes and high levels of quiescence and growth arrest genes when comapared with all other clusters (Figures 6F, 6G). Other CSC genes that have been linked to endometrial cancer (including CD55 and CD44) were also upregulated in Cluster 3 and expressed in a MAPK14-dependent manner (Figure 6H). Most likely therefore, Cluster 3 represents a *MAPK14*-dependent CSC population. The most highly and uniquely expressed gene in Cluster 3 was N-MYC Downstream-Regulated Gene 1 *(NDRG1*) (Figure 6H). We also identified NDRG1 as a *MAPK14*-dependent phosphoprotein in spheroids (Supplementary Figure S5). We used immunofluorescence microscopy to spatially localize NDRG1-expressing (Cluster 3) cells. Figure 6I shows that cells expressing NDRG1 reside in a non-proliferating intermediate zone of spheroids. Cells expressing the stem cell marker CD55 showed a similar distribution pattern in spheroids (Figure S7H). Using immunoblotting, we confirmed that the expression of NDRG1 protein coincided temporally with induction of CD133 (a CSC marker) during spheroid growth and was *MAPK14*-dependent (Figure 6J). We conclude that *MAPK14* is important for sustaining a sub-population of non-proliferating cells that express CSC markers and NDRG1 (a protein implicated in proliferation, metastasis and chemoresistance).

### MAPK14 promotes cancer stemness characteristics

We measured spheroid-forming activity from single cells as an established phenotypic assay for stemness. As shown in Figure 7A-B, spheroid-formation activity of *MAPK14^-/-^* HEC-50B cells was reduced by 0.68-fold when compared with *MAPK14^+/+^* controls. When we seeded equal numbers of *MAPK14^+/+^* or *MAPK14^-/-^* cells into ULA wells to promote multicellular growth, the resulting *MAPK14*-deficient spheroids attained significantly smaller sizes than *MAPK14^+/+^* spheroids (Figure 7C-E). Finally, we tested the effect of *MAPK14* on cancer stemness using a definitive functional assay, namely tumor-seeding activity in a mouse peritoneal dissemination model that recapitulates key features of human disease.^59, 60^ As shown in Figure 7F-G, formation of intraperitoneal tumors was significantly reduced in animals injected with *MAPK14*^-/-^ HEC-50B cells when compared with *MAPK14^+/+^* control groups. We conclude that *MAPK14* is important for stemness and tumorigenicity of HEC-50B cells.

**Figure 7.**
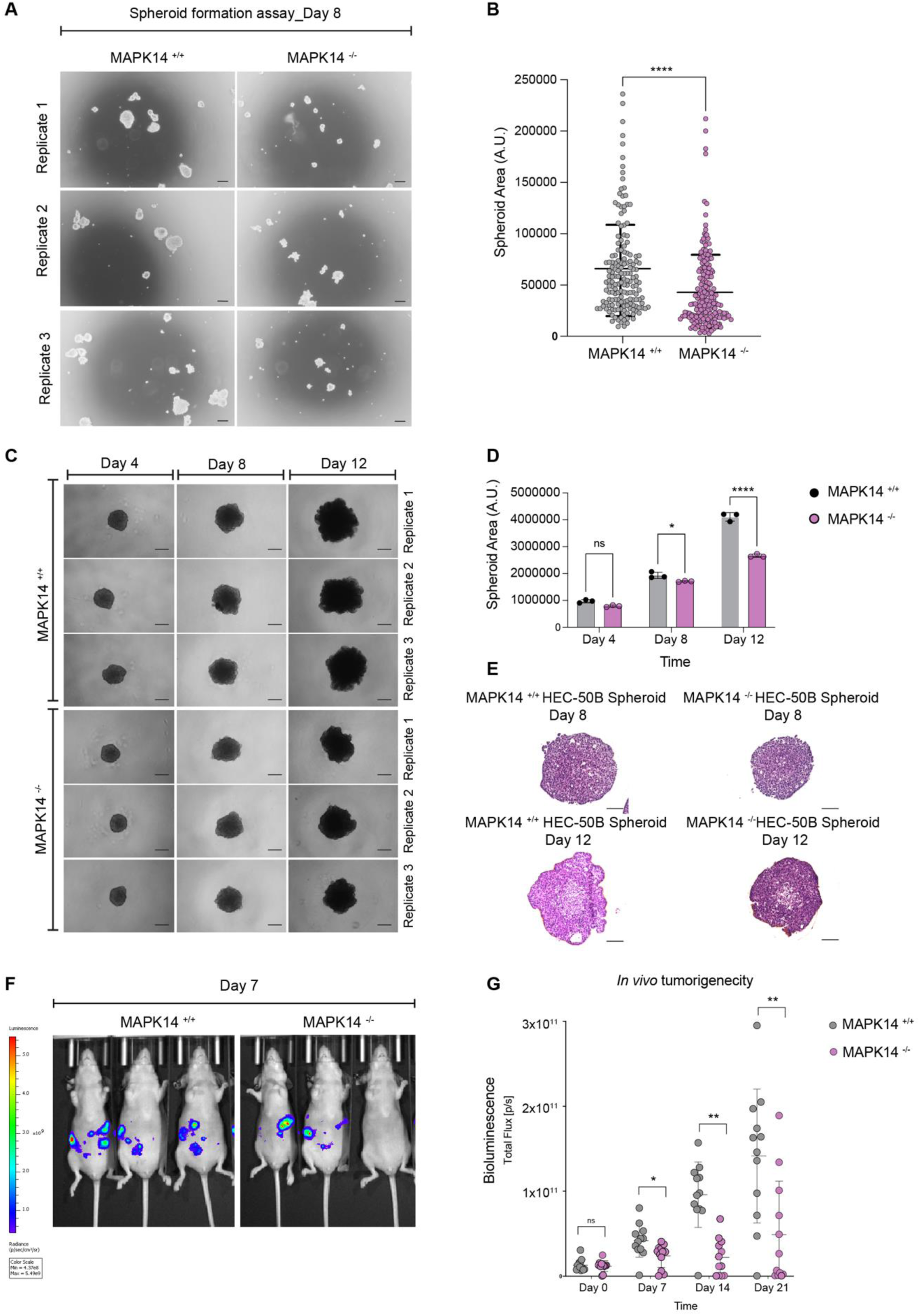
*MAPK14* promotes cancer stemness characteristics. **(A-B)** Representative bright-field images of ULA Plates showing spheroid formation efficiency of *MAPK14^+/+^* and *MAPK14^-/-^* HEC-50B cells (A) and quantification of spheroid areas measured using ImageJ (B). Each dot represents a single spheroid. Graph depicts Mean ± SD. **(C-D)** Representative images of spheroid growth over a 12 day period in *MAPK14^+/+^* and *MAPK14^-/-^* HEC-50B cells/well (C) and quantitative analysis of spheroid areas over time for *MAPK14^+/+^* and *MAPK14^-/-^* HEC-50B cells (D). Each data point on the bar chart depicts Mean ± SD. **(E)** H&E-stained sections of representative *MAPK14^+/+^* and *MAPK14^-/-^* spheroids on days 8 and 12 after seeding. **(F-G)** Representative bioluminescence images of *MAPK14^+/+^* or *MAPK14^-/-^* tumors generated *in vivo* at Day 7 post-intraperitoneal injection of *MAPK14^+/+^* or *MAPK14^-/-^* HEC-50B cells in ovariectomized nude mice (F) and Quantitative analysis of metastatic-tumor growth measured using bioluminescence imaging in mice injected with *MAPK14^+/+^* or *MAPK14^-/-^* HEC-50B cells at indicated timepoints (*n=12*) (G). In (B), unpaired two-tailed t-test was used to determine significance. In (D) Two-way ANOVA was used to infer statistical significance between means of *MAPK14^+/+^* or *MAPK14^-/-^* HEC-50B cells in 3D with time. In (G) unpaired two-tailed t-test was used to determine statistical difference at individual time points between the two comparison groups. ∗p < 0.0332, ∗∗p < 0.0021, and ∗∗∗p < 0.0002, ∗∗∗∗p<0.0001

### HGEC patient tumors contain malignant cell sub-populations resembling MAPK14-dependent CSCs

We interrogated available published scRNASeq profiles of six EC patient tumors to identify EC cells with a gene expression signature corresponding to the HEC-50B spheroid Cluster 3 transcriptome. We identified 12 transcriptionally-discrete clusters (designated ‘Patient Tumor Clusters’ 1-12, Figures 8A, 8B). Interestingly, Patient Tumor Cluster 5 represented a population of cells with a transcriptome matching that of HEC-50B spheroid Cluster 3. Similar to the HEC-50B spheroid Cluster 3, cells in Patient Tumor Cluster 5 expressed high levels of *NDRG1* (Figure 8C) and CSC markers (Figure8D). Cluster 5 cells also expressed low levels of cell cycle genes (Figure 8E) and high levels of growth arrest markers (Figure 8F). GO analysis showed that patient tumor cluster 5 also expressed hypoxia response genes (Figure 8G). Therefore, a *MAPK14*-dependent subpopulation of cells that arises in HEC-50B cells only during spheroid culture has a close counterpart in HGEC patient tumors. We conclude that the unique *MAPK14*-dependent biology we have revealed in our studies with 3D cultures closely recapitulates features of HGEC tumors in patients.

**Figure 8.**
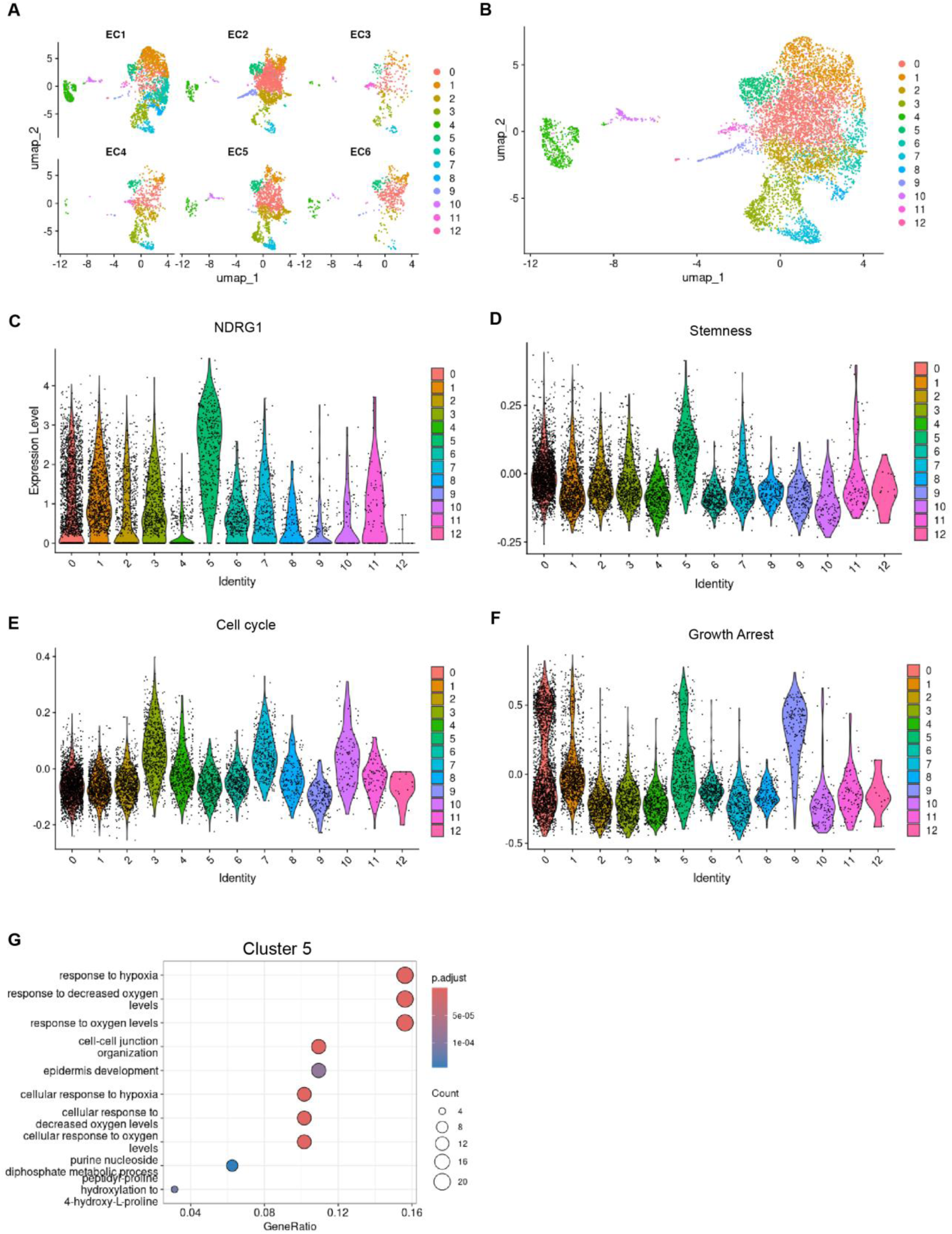
HGEC patient tumors contain malignant cell sub-populations resembling *MAPK14*-dependent CSCs. **(A-B)** UMAP plots of the 13 transcriptionally distinct clusters identified by Seurat in HGEC patient tumors. (A) Separated by patient; (B) Integrated. **(C)-(F)** Plots showing relative levels of *NDRG1* mRNA **(C)** and gene expression signatures for cancer stemness **(D)**, DNA replication and cell cycle **(E)** and quiescence/growth arrest **(F)** in each of the 12 clusters identified by Seurat in scRNA-seq. **(G)** GO enrichment analysis of marker genes expressed by cells in Seurat cluster 5 from endometrial cancer.

## Discussion

Using 3D models we identify the p38α pathway as a key signaling node that is both activated and necessary for diverse tumorigenic programs (including phosphorylation, transcription, and metabolic networks) in spheroids but not in monolayers. Literature on the roles of p38α in growth control and cancer is contradictory and confusing. Although p38α was originally defined as a tumor suppressor,^61^ it was later reported that p38α enhances proliferation^62^ and late-stage tumor progression.^62^ p38α is activated both in response to mitogens and anti-proliferative stresses.^39^ Some studies support a pro-CSC role for p38,^63^ while others report that p38 attenuates CSC properties in skin,^64^ colon,^65, 66^ lung,^67, 68^ brain,^69^ and breast cancers.^70^ Thus there is no consensus on the role of p38α as a pro- or anti-cancer protein, or its involvement in cancer stemness. The remarkable specificity with which p38α drives biological features of endometrial cancer cells only when in 3D culture further illustrates the pleiotropic nature of p38α signaling in different cancer settings.

What then are distal p38α signaling mechanisms that confer spheroid growth and stemness in EC cells? We show that genes and proteins involved in cytokine / chemokine signaling are highly upregulated in spheroids when compared with monolayers and that many of these changes are *MAPK14*-dependent. Our data showing changes in metabolites known to mediate inflammation (PUFAs, cortisol^71, 72^) provide additional support for a role of *MAPK14* in promoting proinflammatory phenotypes of spheroids. Cytokine and chemokine signaling pathways are attractive candidate mediators of p38α-induced proliferation and stemness in endometrial cancer spheroids. Autocrine and paracrine cytokine signaling are implicated in the growth of many cancers including HGEC.^73, 74, 75, 76, 77, 78, 79^ Cytokine networks also promote cancer stemness^76, 80, 81^, providing a possible mechanism for the *MAPK14*-dependent expression of CSC markers in EC spheroids. Moreover, cytokine-dependent inflammation is a hallmark of cancer.^82^ The many cytokines present in tumors are not readily druggable.^83^ However, targeting master regulators of cytokine and chemokine expression (such as p38α) may offer opportunities to block tumor-promoting cytokine signaling.

We identify *NDRG1* as an *MAPK14*-dependent gene whose expression is associated specifically with subpopulations of cells harboring a cancer stemness signature, both in spheroids and in patient tumors. *NDRG1* is inducible by stress, hypoxia, p53 and DNA damage,^49, 50, 51, 52^ but has not previously been linked to stemness. However, NDRG1 interacts with DNA repair enzymes and promotes chemoresistance,^84^ which is a cancer stemness hallmark. Therefore NDRG1 is an attractive candidate mediator of p38α-dependent tumorigenic characteristics in endometrial cancer. It is likely that no single downstream effector mediates all of the biological phenotypes resulting from p38α signaling. By analogy, ERK1-mediated mitogenesis is mediated by numerous downstream substrates and targets that constitute a broad network of integrated biological processes.^85^

Given the important role we have demonstrated for p38α in programming spheroid biology, it will be important to determine whether MK2 and the other protein kinase cascades upregulated in 3D cultures contribute key tumorigenic characteristics in endometrial cancer. From our KSEA, the cyclic GMP (cGMP)-dependent protein Kinase G (PKG), TANK-binding kinase 1 (TBK1), and Ribosomal protein S6 kinase beta-1 (S6K1) were specifically and highly induced during 3D growth. These protein kinase pathways have not previously been linked to 3D spheroid growth or endometrial cancer. PKG1 and PKG iso1 are cGMP-activated protein kinases that promote breast cancer migration and invasion^86^. In our metabolomics profiling studies, cGMP was elevated by 3.2-fold in spheroids when compared with monolayers, consistent with increased activity of the PKG pathway duing 3D growth. TBK1 was initially identified as a kinase responsible for expression of type I interferons (IFNs)^87^ and promotes oncogenic phenotypes of several cancers^88^. Our proteomics and RNASeq experiments show that IFNs and other inflammatory cytokines are highly induced in 3D cultures when compared with 2D monolayers. Cytokines in the tumor microenvironment (TME) promote angiogenesis, epithelial to mesenchymal transition (EMT), invasion and cancer stemness^76, 80, 81^. STING is a downstream TBK1 target and is hyper-phosphorylated in 3D cultures compared with 2D monolayers. Therefore, TBK1-cytokine signaling may also contribute important tumorigenic features of HGEC spheroids. S6K1 is a crucial effector of mTORC1 which directly regulates ribosome biogenesis, cell cycle progression, protein synthesis and metabolic reprogramming^89, 90^. Thus, based on their known functions in other biological contexts, it is likely that the concerted actions of PKG, TBK1, and S6K1 may contribute to endometrial cancer spheroid pathobiology.

If indeed multiple protein kinase pathways contribute to the tumorigenicity of endometrial cancer cells, co-targeting these pathways may achieve cytotoxicity in situations where monotherapies are ineffective. Mice lacking *Pkg1* and *S6k1*^91, 92^ and humans lacking both *TBK1* alleles are viable.^93^ Therefore, if PKG1, S6K1 and TBK1 are important dependencies of endometrial cancer cells, specific therapeutic inhibitors of these kinases will likely lack side-effect toxicities to normal cells.

Our phosphoproteome analyses also revealed considerable reprogramming of the kinome in spheroids lacking *MAPK14*. For example, Aurora kinase B (AURB) and NEK9 were upregulated in *MAPK14^-/-^* spheroids when compared with *MAPK14^+/+^*. AURB functions in the attachment of the mitotic spindle to the centromere^94^ and NEK9 controls centrosome separation.^95^ p38α is active during G2/M^96^ and has a role in satisfying the mitotic checkpoint.^97^ Therefore, in the absence of p38α, mitotic kinases (AURB, NEK9) may compensate to promote G2/M progression. SGK1 was one of the most downregulated kinases in *MAPK14^-/-^* 3D cultures when compared with *MAPK14^+/+^* spheroids. We validated that phosphorylation of NDRG1 at major SGK1 target sites^98^ was *MAPK14*-dependent. In Glioblastoma, SGK1-mediated NDRG1-mediated phosphorylation promotes chemoresistance.^84, 98^ Moreover, SGK1 is a mediator of stress responses to therapeutic agents that promotes tumor progression.^99^ Therefore, the SGK1/NDRG1 pathway may be an attractive candidate for targeted therapy.

Only a limited number of past studies with have identified genes and pathways that are important for 3D growth but dispensable for cells in monolayer culture. A CRISPR screen identified a role for NRF2 in proliferation and survival of A549 lung adenocarcinoma spheroids^55^. An independent study using lung adenocarcinoma cells (A549 and H23) revealed that carboxypeptidase D (which removes a C-terminal RKRR motif from the the insulin-like growth factor 1 receptor α-chain to promote receptor activation) was important for spheroid growth. Both NRF2 activity and carboxypeptidase D expression are associated with poor outcomes for lung cancer patients^55, 100^. However, NRF2 was not activated or required during spheroid growth of breast cancer cells^101^. By analogy, we find that p38α activation during 3D growth is not evident in PDAC spheroids. Therefore, different cancer cell types may deploy distinct mechanisms to sustain multicellular growth. It will be necessary to identify the genes and pathways that define and sustain 3D growth in each unique cancer setting if we are to target those pathways for therapeutic purposes.

Many questions remain regarding how 3D growth creates new molecular programs and dependencies that are absent from conventional 2D cultures. Unlike monolayer cultures, in which cells are relatively homogeneous, isogenic cells growing as multicellular structures such as spheroids exhibit heterogeneity that is driven by a combination of factors including chemical and oxygen gradients,^102^ and mechanical forces^103, 104^. Acidity and hypoxia are potential sources of intrinsic stress and genotoxicity^105, 106, 107, 108^ that could contribute to the unique demands of cells in 3D structures. Further work is necessary to determine how spatial organization and zonation are established in spheroids, and how the molecular programs we identified localize/reside in relation to gradients of cell proliferation, oxygen and pH. Elucidating how cell-cell communication and spheroid architecture are established in a multicellular structure may reveal opportunities to disrupt those interactions and compromise cancer cell viability. Our study demonstrates the enormous impact of pathologically-relevant 3D growth on the biology of EC cells. The model systems, screening platforms and datasets we have generated are resources that will help stimulate future research into mechanisms of EC etiology and therapeutics.

## Acknowledgments

We thank Dr. Pablo Ariel for his assistance in microscopy image acquisition and data analysis. We thank Prof. Shawn Hingtgen for gifting the FLuc-mCh-puro Lentivirus vector. We thank the members of Advance Analytics core (AA core) at Center for Gastrointestinal Biology and Disease, UNC at Chapel Hill for support in single-cell sequencing data acquisition. AA core is supported by CGIBD center grant (P30 DK034987). We thank members of the UNC Lineberger Preclinical Research Unit (PRU) for assistance with *in vivo* experiments which were performed at the University of North Carolina at Chapel Hill and is supported in part by an NCI Center Core Support Grant (CA16086) to the UNC Lineberger Comprehensive Cancer Center. We thank Gabriela De La Cruza and Yongjuan Xia in the Pathology Services Core (PSC) for expert technical assistance with FFPE tissue processing and H &E staining. The PSC is supported in part by an NCI Center Core Support Grant (P30CA016086). National Institutes of Health (R01 ES009558, CA215347, CA229530 to C.V.); University of North Carolina Lineberger Developmental Funding Program Stimulus Award (to J.L.B and C.V.); University of the North Carolina Center for Environmental Health and Susceptibility 2023–2024 Pilot Project Awards (to J.L.B.)

## Author contributions

S.J., J.L.B., and C.V. conceived the study and designed the experiments. S.J., A.M., A.M., and B.R. performed wet-lab experiments. V.B-J. generated PDX models. S.J., X.Z., G.D., B.R., and S.L., analyzed data and prepared figures. D. W., L.H., J.L.B., and C.V. supervised the work. S.J., J.L.B., and C.V. generated the first draft of the manuscript. All authors contributed sections to subsequent drafts and approved the final manuscript.

## RESOURCE AVAILABILITY

### Lead contacts

Further information and requests for resources and reagents should be directed to and will be fulfilled by the lead contacts Cyrus Vaziri and Jessica Bowser.

### Materials Availability

All plasmids and cell lines generated by this study are available upon request from the lead contacts with a completed Materials Transfer Agreement (MTA).

### Data and code availability

TCGA-UCEC datasets were downloaded from TCGA portal (https://portal.gdc.cancer.gov/). The raw data of endometrial cancer patient single cell RNA-seq data were previously reported^109^ and can be found at https://zenodo.org/records/393781. All transcriptomics data analysis (bulk RNA-seq, single-cell RNA-seq, TCGA bulk RNA-seq) were conducted using R statistical language (Version 4.1.0). Bulk mRNA sequencing dataset is available under BioProject ID: PRJNA1120648. Metabolomics analysis is available at the NIH Common Fund’s National Metabolomics Data Repository (NMDR) website, the Metabolomics Workbench, under Project ID: PR002025.

## Method Details

### Cell lines

All endometrial cancer cell lines HEC-50B, HEC-1-A, MFE-296 and HES-UPSC were cultured in RPMI-1640 medium (Gibco, 11875093) supplemented with 10% FBS (Gibco, 26140079) and 1% penicillin/streptomycin (Gibco, 15140122). Other cancer cell lines U-87, A549, FaDu, KYSE-30, Pa02C, Pa03C were cultured in DMEM medium (Corning, 10013CV) supplemented with 10% FBS and 1% penicillin/streptomycin. All cells were cultured at 37°C with 5% CO2. Cell lines were tested for mycoplasma contamination with the Venor®GeM qEP kit and were confirmed negative.

### 3D spheroid and PDX-derived 3D cell culture

For 3D spheroid culture, cells were cultured in ultra-low attachment (ULA) 96-well plates (Corning, 650970) in DMEM/F-12 medium (Gibco, 11330032) supplemented with 20 ng/ml human EGF recombinant protein (Gibco, PHG0314), 20 ng/ml human basic FGF recombinant protein (bFGF) (Gibco, 13256029), 2% B27 supplement (Gibco, 17504044) and 1% penicillin/streptomycin. For multicellular 3D spheroid culture, HEC-50B cells were seeded in ULA 96-well plates, at a seeding density of 2000 cells/well for a duration of 4-12 days. Media was refreshed every two days by aspirating 50% volume and replenishing with fresh media. For passaging 3D spheroids, spheroids were pelleted by centrifugation at *400g* for 5 min. Spheroid pellets were then washed with 1x DPBS (Corning, 21031CV). After aspirating DPBS, spheroids were resuspended in Accutase (Sigma, AT104) at a density of ∼300 spheroids/ml at 37°C for 5 min with intermittent agitation for uniform dissociation into single cells.

Endometrial Cancer Patient Derived Xenografts (PDX) E-J00011615-1 (labelled as PDX-1) and U190801-1 (labelled as PDX-2) were processed for culture in ULA plates as 3D spheroids. Harvested xenograft was washed with 1x cold DPBS. Tissue was re-suspended in cold DMEM/F-12 medium and minced with sterile scalpel. The minced tissue was mechanically dissociated by pipetting and then pressed through 40 µm nylon cell strainer (Falcon, 352340) using a syringe plunger. The remaining tissue fragments on the strainer were resuspended in DMEM/F-12 basal media and digested with 250 U/ml Collagenase Type II (Worthington, 4176) at 37°C water bath for 10 min. Once digested, the enzymatically dissociated cells were pooled with mechanically dissociated cells and was centrifuged at *400g* for 5 min. The cell pellet was resuspended in room temperature DPBS and centrifuged at *400g* for 5 min. This was repeated twice. The final cell pellet was resuspended in DMEM/F-12 medium supplemented with 20 ng/ml EGF, 20 ng/ml bFGF and 1% penicillin and streptomycin and 50 µg/ml Gentamicin (Gibco, 15750060). PDX derived cells were plated in 6-well ULA plates (Corning, 3471). Cells were supplemented with fresh media every 48-72 hours. PDX cultured as 3D spheroids were separated from dead cells and debris by reverse filtration using 40 µm nylon cell strainer. Spheroids were collected on Day 14 of culture for protein extraction.

### CRISPR-Cas9 screen

A pooled sgRNA library was custom designed and synthesized by CustomArray Inc. The library was synthesized to target 504 DNA Damage Repair and stress-response genes. For each gene targeted, 10 sgRNAs targeting functional domains were included. Additionally, 1000 non-targeting (Non-Tgt) sgRNAs were included in the library. Cloning of sgRNAs into LentiCRISPRv2-DD-Cas9 BSD backbone and lentivirus preparation was performed as described previously.^41, 110^ HEC-50B cells cultured as monolayers were transduced with pooled CRISPR-DD-Cas9-BSD-lentivirus at an MOI of ∼0.3 and at a coverage of 500 cells per sgRNA. Transduced cells were selected with Blasticidin (Gibco, A1113903) at a concentration of 10 µg/ml for 5 days. After complete selection, cells were treated with Shield-1 (Takara, 632188) at a concentration of 1 µM for 4 days. Upon complete Shield-1 treatment, ∼6×106 cells were collected as population doubling 0 (PD0) samples for genomic DNA (gDNA) isolation. From the remaining cells, 3x 106 cells were seeded as 2D monolayers and as multicellular 3D spheroids, each. Cells cultured in 2D were passaged after every ∼4 population doublings (3-4 days). Cells cultured in 3D were passaged after every ∼4 population doublings (7 days). After 10 and 20 population doublings (PD10 and PD20) in culture, the 2D and 3D cultured cells were collected for gDNA isolation. gDNA from the PD0, PD10 and PD20 of 2D and 3D cultured cells were isolated using the (QIAGEN, 69506). The sgRNA sequences were amplified using a nested PCR amplification protocol and purified by magnetic beads as described previously (PMID: 38438348). Purified PCR products were pooled 1:1 and a 5% PhiX DNA spike-in was included and sequenced by Illumina HiSeq PE150 platform at Novogene. Quality control, mapping, normalization, and analysis of sequencing data was performed using VOLUNDR pipeline.^111^

### Lentiviral vectors and lentivirus preparation

For validation of CRISPR-Cas9 screen, sgRNA sequences were cloned in LentiCRISPRv2-DD-Cas9-BSD backbone by Hi-Fi assembly as described previously.^41^ All cloned sgRNA constructs were sequence verified. For generation of GFP expressing cell lines, FU-H2B-GFP-IRES-Puro construct (Addgene, 69550) was used. For generation of luciferase expressing cell line, FLuc-mCh-puro Lentivirus vector^112^ was a gift from Dr. Shawn Hingtgen. For generation of lentiviruses, HEK293T cells were co-transfected with 12 µg lentivirus expression plasmid and 4.5 µg each of the packaging and envelope plasmids psPAX2 and pMD2.G in a 10 cm dish with Lipofectamine 2000 (Invitrogen, 11668019) in Opti-MEM medium (Gibco, 31985070). Medium was changed after 6 hours of transfection. Lentivirus containing media was harvested at 24 hours and at 48 hours, pooled, and filtered through 0.45 µm membrane (Corning, 431225) prior to infecting target cells.

### Generation of stable cell lines

For viral transduction, target cells were incubated overnight with lentivirus-containing medium with 8 µg/ml polybrene (Sigma, TR-1003-G). After 48 hours, infected cells were trypsinized and replated in growth medium containing the appropriate selection antibiotic. For HEC-50B cells transduced with mCherry-luciferase encoding lentivirus, selection was performed with 1 µg/ml of Puromycin (Gibco, A11138-03) for 3-4 days. For HEC-50B cells transduced with H2B-GFP encoding lentivirus, selection was performed with 1 µg/ml of Puromycin for 3-4 days. For HEC-50B cells transduced with sgRNA-Cas9 encoding lentivirus, selection was performed with 10 µg/ml Blasticidin for 5 days. For stabilization of DD-Cas9 and initiating genome editing, cells were treated with 1 µM Shield-1 for 4-7 days.

### Competitive growth assay for CRISPR screen validation

For 2D competitive growth assay, cells expressing gene-targeting sgRNA (sgRNA-Tgt) or non-targeting sgRNA (sgRNA-Non-Tgt), were mixed with cells expressing H2B-GFP in 1:1 ratio and seeded in 6-well plates. Cells in 2D culture were passaged every 3-4 days. For competitive growth assays in spheroids, sgRNA-Tgt or sgRNA-Non-Tgt expressing cells were mixed with H2B-GFP expressing cells in 1:1 ratio, and seeded in 96-ULA plates. Cells in spheroid culture were passaged every 7 days. Media changes were performed every 48 hours by aspirating 50% volume of media and replenishing with fresh media. At every passage, cells in monolayer or spheroid culture were dissociated and analyzed by flow cytometry to determine numbers of GFP-positive and GFP-negative. Rations of of GFP-positive and GFP-negative cells were quantified to analyze relative growth. All comparisons were performed using triplicate cultures.

### Cell cycle and cell death analysis of 3D spheroids

Cell cycle profiles and cell death measurements were performed by flow cytometric analysis. For cell cycle profile analysis, HEC-50B cells were cultured as 3D spheroids for 12 days, then incubated with 10 µM BrdU for 3 hours. BrdU-labeled spheroids were collected and dissociated with Accutase. Dissociated cells were fixed in cold 65% Ethanol in DMEM at 4°C overnight, then stained with FITC-conjugated anti-BrdU antibody and propidium iodide (PI) and analyzed by flow cytometry as described before.^41^ For cell death analysis of HEC-50B 3D spheroids cultured for 12 days, then dissociated using Accutase. Unfixed dissociated single-cells were labeled with Annexin V antibody to detect apoptotic cells. Necrotic cells were labelled with PI. Assays were performed using BD Pharmingen™ FITC and PE Annexin V Apoptosis Detection Kit (BD, 556547) according to the manufacturer’s instructions. The percentage of live and dead cells were analyzed by flow cytometry. All flow cytometric analysis were performed using Accuri C6 plus flow cytometer (BD) and analyzed using FlowJo 10.8.2 software.

### Immunoblot analysis

For whole cell extracts, monolayers or spheroids were collected and pelleted at *400g* for 5 min, washed with cold 1x PBS and pelleted again by centrifugation. The cell pellets were resuspended in CSK buffer (10 mM Pipes, pH 6.8, 100 mM NaCl, 300 mM sucrose, 3 mM MgCl_2_, 1 mM EGTA, 1 mM dithiothreitol, 0.1 mM ATP, 1 mM Na_3_VO_4_, 10 mM NaF and 0.1% Triton X-100) with freshly added protease (Roche, 4693159001) and phosphatase inhibitors (Roche, 4906837001) and incubated at 4°C for 20 min on a rotator. Nuclease (Pierce, 88702) was added to the lysate, followed by 10 min incubation at RT. Samples were then re-incubated on ice for 10 min and then sonicated with a 5s pulse at 20% amplitude. Lysates were then centrifuged at *20,000g* for 10 min at 4°C to remove cellular debris. Supernatant was collected as whole cell extracts. Protein concentration was measured by Bradford assay (Biorad, 5000006). Equal amounts of protein lysates are denatured in Laemmli SDS buffer and were run on 4–20% Tris-Glycine gels (Invitrogen, XP04205BOX), and transferred to nitrocellulose membrane. Membranes were blocked with 5% w/v non-fat dry milk in Tris-buffered saline with 0.1% Tween-20, then incubated with various primary antibodies in the same buffer. Bound primary antibodies were detected using either an anti-rabbit or an anti-mouse HRP-conjugated secondary antibody, at 1:5000 dilution, in %5 w/v milk. HRP-conjugated antibodies were detected by chemiluminescence using Revvity Western Lightning Plus ECL (NEL104001EA) and developed on autoradiography films (Genesee, 30-810).

### EPO-luciferase activity

EPO-HRE-luciferase plasmid was used as a reporter of HIF1α activity. pGL2-luciferase plasmid was used as a promoterless control. 60 ng EPO-HRE-expression plasmid and 60 ng pRenilla-plasmid were co-transfected into HEC-50B cells plated in 24-well F-bottom plates. At 24 hours post transfection, cells were collected and seeded as 2D monolayers in 96-well F-bottom plates or as 3D spheroids in ULA-96-well-U-bottom plates. 48 hours later, cells were harvested, and luciferase activities were measured by using the dual luciferase reporter assay (Promega, Madison, WI, USA) according to the manufacturer’s protocol. For each experimental condition, transfections were performed in triplicate.

### Mitochondrial superoxide detection assay

HEC-50B cells were cultured as monolayers and as 3D spheroids for a duration of 8 days. Then, cells were collected, dissociated with Accutase and analyzed using MitoSox Green Superoxide detection assay kit (Invitrogen, M36006) according to the manufacturer’s protocol. All assays were performed in triplicate. Samples were analyzed by flow cytometry and data was analyzed using FlowJo analysis software.

### Quantitative RT-PCR analysis

HEC-50B cells were cultured as 2D monolayers and 3D spheroids for a duration of 8 days. Cells were washed twice with 1X DPBS twice, then directly collected in RNA lysis buffer and RNA isolation was performed according to the manufacturer’s protocol using RNeasy Plus Mini Kit (Qiagen, 74106). cDNA synthesis was performed using iScript™ cDNA Synthesis Kit (Biorad, 1708891) as per manufacturer’s instructions. RT-PCR analysis was performed using iTaq Universal SYBR Green Supermix (Biorad, 1725121) on Applied Biosystems 7500 Fast Real-Time PCR System. For quantification of gene expression, the 2–ΔΔCt method was used. GAPDH expression was used for normalization. All real-time PCR primers are listed in Supplementary Table X.

### Transcriptomic analysis

For transcriptomic analysis, *MAPK14^+/+^* and *MAPK14^-/-^* triplicate cultures of HEC-50B cells were grown as monolayers and as 3D spheroids for 8 days. RNA was extracted from cells using the RNeasy Plus Mini Kit according to manufacturer’s protocol. RNA sample QC and mRNA library preparation was performed by Novogene and the Illumina NovaSeq 6000 PE150 platform was used for sequencing. For bulk mRNA-seq analysis, R package DESeq2 (Version 1.32.0)^113^ was used to normalize the gene count data and perform the differential expression analysis. For the differential expression analysis, the gene count data were fitted to the Negative Binomial distribution and the Wald test was used for testing the difference between a full and reduced model. Genes with adjusted p-value < 0.1 and log2 fold change < −1 or > 1. After acquiring the differential expressed genes (DEGs), R package ClusterProfiler (Version 4.0.5)^114^ was used for Gene Ontology (GO) enrichment analysis. Pathways with adjusted p-value < 0.05 in the Over Representative Analysis (ORA) were considered statistically significant. p-values were corrected by Benjamini and Hochberg method for multiple comparison. For visualization of the specific pathway genes of bulk RNA-seq data across different genotype and culture, R package ComplexHeatmap(Version: 2.8.0)^115^ was used for generating the heatmaps from normalized count data

### Mass spectrometry analysis of the HEC-50B proteome and phosphoproteome

To prepare samples for mass spectrometry, *MAPK14^+/+^* and *MAPK14^-/-^* HEC-50B cells were cultured as monolayers and as 3D spheroids for 12 days. Whole cell extracts from monolayers or spheroids were prepared as described in immunoblot analysis section. All sample groups included biological triplicates. Cell lysates (600 µg; n=3) were lysed in CSK lysis buffer (10 mM Pipes, pH 7, 100 mM NaCl, 300 mM Sucrose, 3 mM MgCl2, 1mM EGTA, 10 mM NaF, 25 mM β-glycerophosphate, and 0.1% Triton X-100; freshly supplemented with cOmplete Protease Inhibitor Cocktail (Roche) and PhosSTOP inhibitors (Roche), reduced with 5mM DTT for 45 min at 37°C and alkylated with 15mM iodoacetamide for 45 min in the dark at room temperature. Samples were digested with LysC (Wako, 1:50 w/w) for 2 hr at 37°C, then diluted to 1M urea and digested with trypsin (Promega, 1:50 w/w) overnight at 37°C. The resulting peptide samples were acidified to 0.5% trifluoracetic acid, desalted using desalting spin columns (Thermo), and the eluates were dried via vacuum centrifugation. Peptide concentration was determined using Quantitative Colorimetric Peptide Assay (Pierce).

Samples were labeled with TMTpro 16plex reagents (Thermo Fisher).150 µg of each sample was reconstituted with 50 mM HEPES pH 8.5, then individually labeled with 400 µg of TMT reagent for 2 hr at room temperature shaking in a Thermomixer. Prior to quenching, the labeling efficiency was evaluated by LC-MS/MS analysis of a pooled sample consisting of 1ul of each sample. After confirming >98% efficiency, samples were quenched with 50% hydroxylamine to a final concentration of 0.4%. Labeled peptide samples were combined 1:1, desalted using Thermo desalting spin column, and dried via lyophilization. The dried TMT-labeled sample was fractionated using high pH reversed phase HPLC^116^. Briefly, the samples were offline fractionated over a 90 min run, into 96 fractions by high pH reverse-phase HPLC (Agilent 1260) using an Agilent Zorbax 300 Extend-C18 column (3.5-µm, 4.6 × 250 mm) with mobile phase A containing 4.5 mM ammonium formate (pH 10) in 2% (vol/vol) LC-MS grade acetonitrile, and mobile phase B containing 4.5 mM ammonium formate (pH 10) in 90% (vol/vol) LC-MS grade acetonitrile. The 96 resulting fractions were then concatenated in a non-continuous manner into 24 fractions and 5% of each were aliquoted for total proteome analysis, lyophilized and stored at - 80°C until further analysis. The remaining 95% of each fraction was further concatenated into 4 fractions and dried down via vacuum centrifugation. For each fraction, phosphopeptides were enriched with the High Select Fe-NTA kit (Thermo) per manufacturer’s protocol. The Fe-NTA eluates were lyophilized and stored at −80°C until further analysis.

The proteomics samples consisting of 24 fractions for the proteome analysis and 6 fractions for the phosphoproteome analysis were analyzed by LC/MS/MS using an Ultimate 3000-Exploris480 (Thermo Scientific). Samples were injected onto an IonOpticks Aurora series 2 C18 column (75 μm id × 15 cm, 1.6 μm particle size), and the proteomics samples were separated over a 70 min method, while a 100 min method was used for the phosphoproteomics samples. The gradient for separation consisted of 5–45% mobile phase B at a 250 nl/min flow rate, where mobile phase A was 0.1% formic acid in water and mobile phase B consisted of 0.1% formic acid in 80% acetonitrile. The Exploris480 was operated in turboTMTpro mode with a cycle time of 3s. Resolution for the precursor scan (m/z 375–1400) was set to 60,000 with an AGC target set to 100%; maximum injection time set to auto Resolution of the MS2 scan was set to 30,000, and consisted of HCD normalized collision energy (NCE) 33 for the proteomics samples and 34 for the phosphoproteomics samples; AGC target set to 300%; maximum injection time of 55 ms; isolation window of 0.7 Da.

Raw data files were processed using Proteome Discoverer (PD v2.5, Thermo Fisher), set to ‘reporter ion MS2’ with ‘16pex TMT’. Peak lists were searched against a reviewed Uniprot human database (downloaded January 2023 containing 20,401 sequences), appended with a common contaminants database, using Sequest HT within Proteome Discoverer. Data were searched with up to two missed trypsin cleavage sites, fixed modifications: TMT16plex peptide N-terminus and Lys, carbamidomethylation Cys, dynamic modification: N-terminal protein acetyl, oxidation Met. Precursor mass tolerance of 10 ppm and fragment mass tolerance of 0.02 Da (MS2). Peptide false discovery rate was set to 1%. Reporter abundance was calculated based on intensity. Razor and unique peptides were used for quantitation.

PSM data was exported from PD and used as input for our in-house data analysis script which uses the MSstatsTMT package^117^ for protein summarization and statistical analysis in R [V 4.3.0 R Core Team (2021). R: A language and environment for statistical computing. R Foundation for Statistical Computing, Vienna, Austria. URL https://www.R-project.org/.]. The following arguments were set to True in the PDtoMSstatsTMTFormat function for the proteomics dataset, useNumProteinColumn, useUniquePeptide, rmPSM_withfewMea_withinRun, removeFreMeasurements. These arguments correspond to removing shared peptides, removing peptides assigned to more than one protein, and removing features with less than 3 measurements within each run, respectively. The same arguments were used for the phosphopeptides data with the exception of setting useNumProteinsColumn to FALSE. For the phosphoproteomics data we used the MSstatsPTM package^118^ for phosphopeptide summarization as well as performing a protein adjusted phosphopeptides normalization. Both datasets were normalized using the ‘msstats’ method and imputed using the dataSummarizationPTM_TMT function.

Kinase-Substrate enrichment analysis was performed in R using a modified version of the KSEAapp package^119^. The kinase-substrate database used was downloaded from phosphositeplus in April of 2023, and was filtered for only human substrates. A minimum of 3 substates were required to keep a kinase.

Phosphoproteome data was uploaded to Ingenuity Pathway Analysis (Qiagen, 2023) with the following cutoffs: p-value < 0.05 and Log2 FC > 0.5 or < −0.5 for each comparison. The canonical signaling pathways affected by the phosphorylated proteins were manually curated to retain only the significantly overrepresented (p-value < 0.05) and significantly activated (z-score > 2) or inhibited (z-score < −2) pathways of relevance.

### Metabolomics Analysis

*MAPK14^+/+^* and *MAPK14^-/-^* HEC-50B cells were cultured as monolayers and as 3D spheroids for 12 days. Cells were washed three times with ice cold PBS, then quenched with 1 volume of −20°C cold Acetonitrile (Fisher, A955-1) and collected with additional 0.75 volume of ultra-pure H_2_O (Pierce, PI51140). All sample groups included biological triplicates. Spheroid extracts were dried by speed vac overnight and then reconstituted in a volume of 95:5 water:methanol that was proportional to each sample’s protein concentration. Samples were vortexed for 10 min at 5000 rpm and then centrifuged for 10 min at 16000 x g at 4°C. Supernatants were transferred to autosampler vials and an aliquot of 10 μL was taken from each sample and combined into a single mixture to make a quality control study pool (QCSP). LC-MS grade water was processed in an identical manner as the study samples to prepare method blanks. An injection volume of 5 μL was used for untargeted analysis.

Metabolomics data were acquired on a Vanquish UHPLC system coupled to a Q Exactive™ HF-X Hybrid Quadrupole-Orbitrap Mass Spectrometer (Thermo Fisher Scientific, San Jose, CA). An HSS T3 C18 column (2.1 × 100 mm, 1.7 µm, Waters Corporation) at 50 °C with binary mobile phase of water (A) and methanol (B), each containing 0.1% formic acid (v/v) was used for metabolite spearation. The mobile phase gradient started from 2% B, and increased to 100% B in 16 min, then held for 4 min, with the flow rate at 400 µL/min. Positive UHPLC-HRMS data was acquired in a mass range from 70 to 1050 m/z under a data-dependent acquisition mode fragmenting the top 20 most abundant ions per scan to collect MS/MS data. Quality control materials (QCSPs and blanks) were interspersed amongst the study samples during data acquisition. Raw data was preprocessed using Progenesis QI (version 2.1, Waters Corporation) for peak picking and alignment. To remove background, peaks with a higher average abundance in blanks compared to QCSPs were removed. Data was normalized to a reference QCSP sample using the “normalize to all” function in Progenesis QI ^120^. Peaks were matched to an in-house standard library and public databases (NIST, HMDB)^121^. Peaks were matched to metabolites by retention time (RT ± 0.5 min, in-house matches only), exact mass (MS, <5 ppm), and MS/MS similarity (>30%). To communicate matching evidence an ontology system was used as described previously^122, 123^. Multivariate analysis was performed using SIMCA 16 (Sartorius Stedim Data Analytics AB, Umeå, Sweden) using the normalized, filtered data. Heatmaps and pathway analyses were performed using MetaboAnalyst 5.0^124^.

### Whole mount live spheroid imaging

Multicellular spheroids were imaged at Day 8 of spheroid growth. Spheroids were incubated with Hoechst 33342 (Invitrogen, R37605) diluted 100x for 20 min. Propidium iodide was added to wells at a final concentration of 8 µg/ml before imaging. Images were acquired using 10x lens of Keyence BZ-X800 Microscope. Images acquired as a z-stack were processed using Keyence imaging software.

### 3D spheroid immunofluorescence analysis

For immunofluorescence staining MAPK14 ^+/+^ or MAPK14 ^-/-^ HEC-50B cells were cultured for 4-12 days as 3D spheroids. To label replicating cells, 10 µM EdU was added to the spheroid culture medium for 6 hours. The EdU-labeled spheroids were collected and washed with 1x DPBS, then fixed with 2% PFA overnight at 4°C on a rotator. After fixation, spheroids were washed with 1X PBS and incubated in 15% sucrose in PBS for 8 hours followed by incubation in 30% sucrose solution in PBS for 16 hours at 4°C. ∼192 spheroids were then pelleted and resuspended in 60 µl 1x PBS and added to intermediate size molds pre-layered with O.C.T medium (Tissue-Tek, 62550) for cryofreezing. Cryofrozen OCT blocks were sectioned into 10 µm thick slices onto Superfrost plus slides (Fisherbrand, 1255015) and stored at −80 °C until further analysis. For immunostaining, frozen slides were thawed to RT for 5 min and washed twice with 1x PBS for 5 min. Incorporated EdU was detected using a Click-iT™ Plus EdU Cell Proliferation Kit for Imaging (Invitrogen, C10640) according to the manufacturer’s instructions. The sections were permeabilized with 0.3% (v/v) Triton X-100 in DPBS for 30 min, then blocked in a solution containing 5% (w/v) bovine serum albumin, 1% (v/v) goat serum in DPBS and 0.1% Triton X-100 at RT for 1 hour. For antibody staining, sections were labelled with a primary antibody solution (containing 5% (w/v) bovine serum albumin, 1% (v/v) goat serum in DPBS and 0.1% Triton X-100) overnight at 4°C in a humidified chamber. The following primary antibodies were used for immunofluorescence staining: cleaved Caspase 3 (CST, 9664S), phosho-H2AX-S139 (Sigma, 05-636), NDRG1 (CST, 9485S) and CD55 (SCBT, sc-51733). After primary antibody incubations, sections were washed three times with 1X PBS for 5 min. The sections were then stained with secondary antibody at a dilution of 1:200 at RT in a humidified chamber. Mounting media containing DAPI was applied to sections (Invitrogen, P36941). Images were acquired using a Zeiss LSM 900 laser scanning confocal microscope. Images were acquired using a 20x lens as Z-stacks and image tiles were stitched together to obtain the final images.

### Spheroid formation assay and multicellular tumor spheroid growth assay

For spheroid formation assays, *MAPK14^+/+^* or *MAPK14^-/-^* HEC-50B cells were seeded as single-cells in suspension in ULA 6-well plates, at a seeding density of 2000 cells/well (6 replicates per experimental condition). Images of spheroids were acquired at Day 8 using a 4x phase contrast lens of Olympus CKX41 inverted Phase Contrast Fluorescence microscope. For each replicate, images were acquired in quadruplicate. The size of spheroids was calculated by ROI area measurements.

To measure growth as multicellular spheroids, *MAPK14^+/+^* or *MAPK14^-/-^* HEC-50B cells were seeded in ULA 96-well plates (2000 cells/ well). Images of spheroids were acquired, at Days 4, 8 and 12 post-seeding using a 4x phase contrast lens. Images were analyzed using ImageJ2 V2.9 software to define ROIs and determine spheroid areas.

### *In Vivo* Tumorigenicity assay

Animal experiments were approved by the Institutional Animal Care and Use Committees at UNC and performed according to guidelines. For *in vivo* tumorigenicity assay, 5-week-old female NU-Foxn^nu^ mice underwent bilateral ovariectomy. For post-operative pain management, mice were injected with 0.1 mg/kg Buprenorphine; Lidocaine every 8-12 hours, as needed. 5 weeks after ovariectomy, mice were injected with *MAPK14^+/+^* or *MAPK14^-/-^* HEC-50B-mCherry-Luciferase expressing cells, intraperitoneally, at a concentration of 2 x 10^6^ cells/100µl in sterile DPBS (*n=12*). Tumor growth was monitored via bioluminescence measurement using 150 mg/kg D-Luciferin 15 min prior to imaging, and images were acquired of ventral whole mice body.

Bioluminescent images were acquired using IVIS-Lumina (PerkinElmer Inc) imager at Day 0 after injection and thereafter weekly. Images acquired were analyzed using Living Image® 4.7.4 Software.

### Single-cell RNAseq analysis of HEC-50B spheroids

*MAPK14^+/+^* and *MAPK14^-/-^* HEC-50B cells were cultured as multicellular 3D spheroids for 12 days. ∼96 spheroids were pooled and dissociated with Accutase into a single-cell suspension. The two sample groups were collected in triplicate.

For single-cell capture, library preparation and sequencing: Cell suspensions were labelled with TotalSeq-B hashtag antibodies (BioLegend) following a modified version of the manufacturer’s sample multiplexing protocol (https://tinyurl.com/ycy5wah8). First, 1.5 μL of hashtag antibody was added to 150 μL of cells per sample and incubated on ice for 30 minutes. Cells were then collected by centrifugation and washed three times with 1.5 mL ice-cold 1X PBS (calcium and magnesium-free). Finally, cells were filtered through a 40 μm cell strainer and stained with the viability markers Annexin V (BioLegend #640918) and SYTOX Blue (ThermoFisher #S34857) following manufacturer’s instructions. Single, live cells were then collected using a Sony SH800 cell sorter. Each hashtag-labeled, sorted sample was counted using the Luna-FL acridine orange/propidium iodide viability fluorescence assay (Logos Biosystems #F23001). Equal numbers of live cells were pooled from each sample, pelleted, and resuspended in 1X PBS (calcium and magnesium-free) + 0.04% BSA. The pooled sample was counted and then prepared for single cell capture, reverse transcription, and library preparation using the Chromium Next GEM Single Cell 3’ v3.1 kit with Feature Barcoding for Cell Surface Protein according to the manufacturer’s user guide (https://tinyurl.com/5n9a93h9). The cell pool was loaded onto three lanes of the Chromium Next Gem Chip G targeting a total recovery of 30,000 cells (5,000 cells per sample). The libraries from each inlet were pooled and sequenced on NovaSeq 6000 S2 flow cell (Illumina) to a depth of ∼77,093 read pairs per cell.

Single-cell RNA-seq data processing and quality control: Read processing was performed using the 10x Genomics workflow. Briefly, the Cell Ranger v3.0.1 Single-Cell Software Suite was used for demultiplexing, barcode assignment, and unique molecular identifier (UMI) quantification http://software.10xgenomics.com/single-cell/overview/welcome). Sequencing reads were aligned to the HG38 reference genome (Genome Reference Consortium Human Build 38) using a pre-built annotation package obtained from the 10X Genomics website (https://support.10xgenomics.com/single-cellgeneexpression/software/pipelines/latest/advanced/references)

### Single-cell RNAseq data analysis

R package Seurat (Version 4.0)^125^ was used for analyzing the single-cell RNA-seq (scRNA-seq) data. For quality control, only cells with more than 200 features and less than 25% mitochondria counts were retained. We used the standard Seurat pipeline to normalize and scale the data then performed the dimensional reduction with linear method PCA and non-linear method UMAP to visualize the data. The cells were clustered using the KNN-graph-based method using the first 30 dimensions and the resolution parameter 0.5 by the function FindNeighbors and FindClusters in Seurat. After acquiring the cluster information, function FindAllMarker was used to find all the marker genes of each cluster with the threshold at min.pct > 0.1 and logfc.threshold > 0.25. Only marker genes with the average log2 fold change > 1 adjusted p-value < 0.05 in Wilcoxon Rank Sum test were considered statistically significant. To obtain pathway expression scores, function AddModuleScore was used by calculating the relative average expression of genes in the pathway and subtracting the aggregated expression of control features. For endometrial cancer patient scRNA-seq data, canonical correlation analysis (CCA) was used to remove the biological differences from different patients and integrate into a single dataset for downstream analysis.

