## Supplementary Figure 1 for "*MAPK14*/p38α Shapes the Molecular Landscape of Endometrial Cancer and promotes Tumorigenic Characteristics"

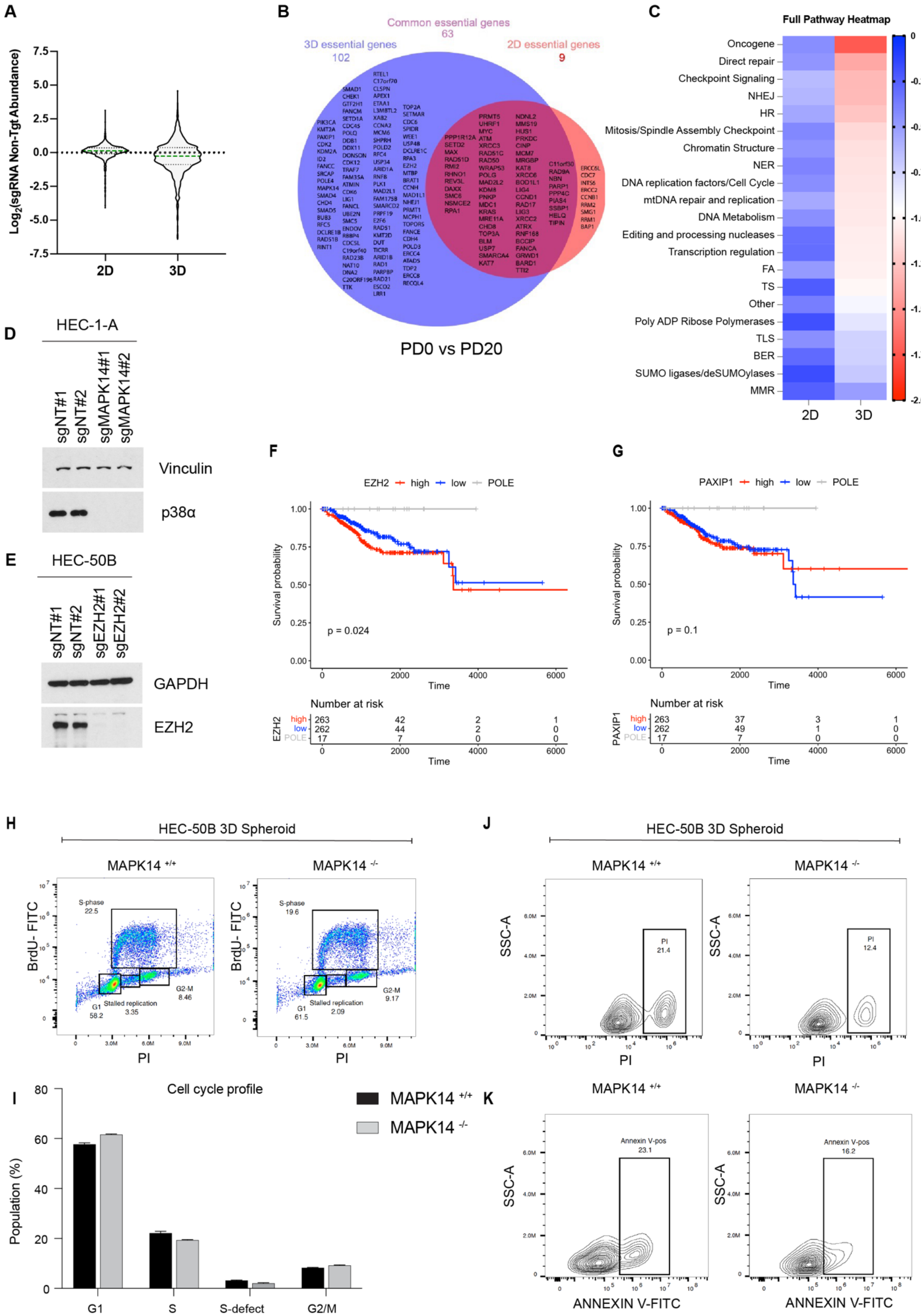

**Supplementary Figure 1. CRISPR dropout screening identifies genes required for growth of HEC-50B cells in spheroid and monolayer culture.**

**(A)** Violin plot depicting abundance of non-targeting (control) sgRNA in HEC-50B cells after 20 population doublings in 2D and 3D culture.

**(B)** Venn diagram showing sgRNAs significantly depleted in spheroid (3D, Blue) and monolayer (2D, Red) cultures. The overlap indicates sgRNAs depleted during both spheroid and monolayer growth.

**(C)** Heatmap showing pathway dependencies of HEC-50B cells growing in monolayer (2D) and spheroid (3D) culture, as determined by relative dropout of sgRNAs under each culture condition.

**(D)** Immunoblot confirming ablation of p38$\alpha$ expression by two independent sgRNAs targeting *MAPK14* in HEC-1-A cell line.

**(E)** Immunoblot confirming ablation of EZH2 expression by two independent sgRNAs in HEC-50B cells.

**(F-G)** Kaplan-Meier curves showing overall survival probability of endometrial cancer patients expressing high (upper quartile) or low (lower quartile) levels of *EZH2* (F) and *PAXIP1* (G) analyzed using patient data from the entire TCGA uterine cancer cohort. Log-rank tests were used to infer statistical significance between high and low gene expression groups.

**(H-I)** Density pseudocolour scatter plots showing cell cycle profiles of $MAPK14^{+/+}$ (left) and $MAPK14^{-/-}$ (right) HEC-50B spheroids. Panel (I) shows quantitative analysis of results from (H) and indicates the percentage of cells in G1, S, and G2/M phases of the cell cycle in $MAPK14^{+/+}$ and $MAPK14^{-/-}$ HEC-50B spheroids. The bars labeled 'S-defects' indicate the numbers of BrdU-negative cells with an S-phase DNA content and represent cells experiencing DNA replication stress.

**(J-K)** Contour plots from flow cytometry experiments indicating the percentage of propidium iodide (PI)-positive cells (J) and the percentage of Annexin-V-FITC-positive cells (K) in $MAPK14^{+/+}$ (left) and $MAPK14^{-/-}$ (right) HEC-50B spheroids

### Supplementary Figure 2

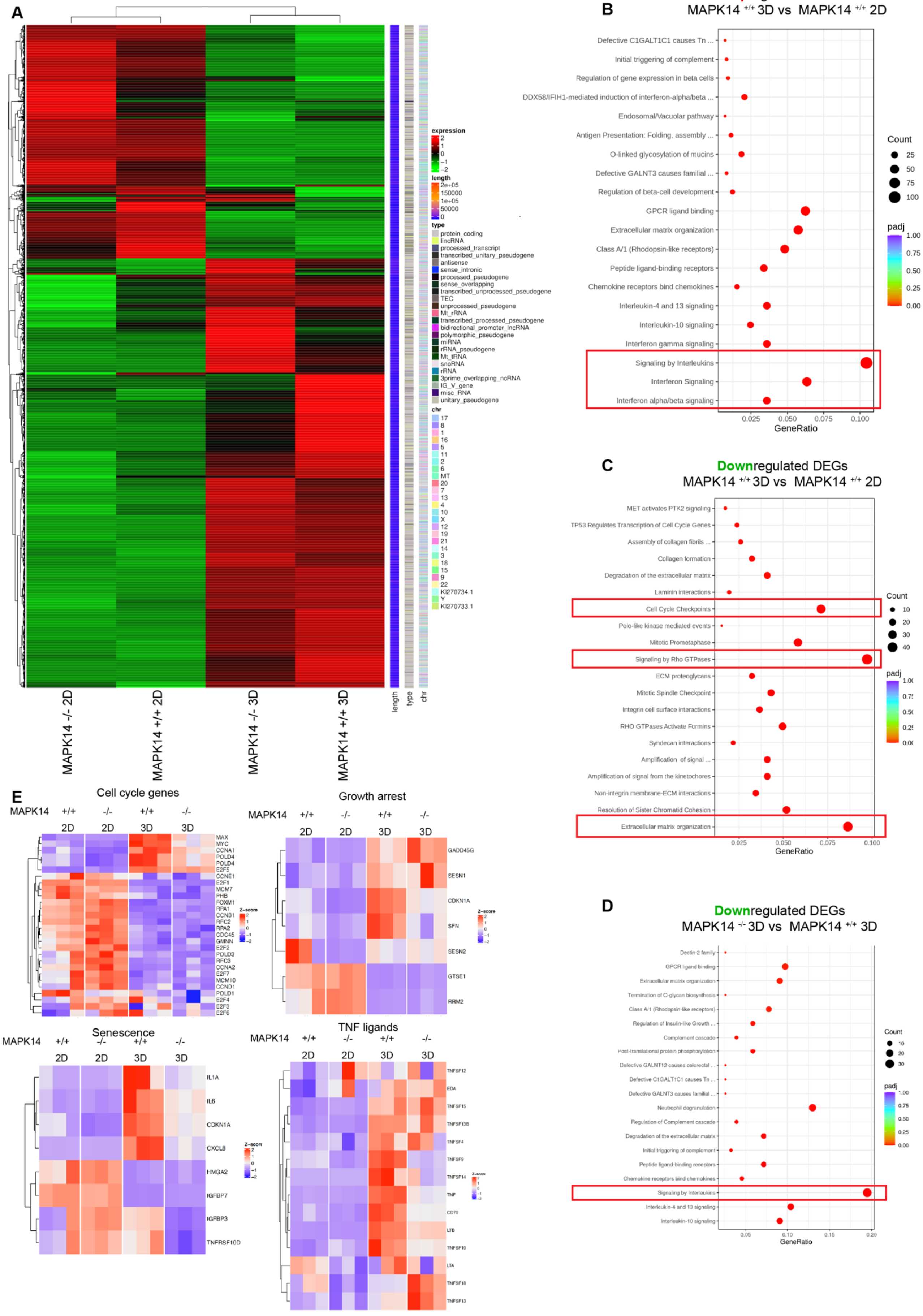

**Supplementary Figure 2. Effect of *MAPK14* on the transcriptome of HEC-50B cells in monolayer and spheroid culture**

**(A)** Heatmap showing differential hierarchical clustering of mRNAs between *MAPK14*$^{+/+}$ and *MAPK14*$^{-/-}$ HEC-50B cells cultured as monolayers and spheroids.

**(B-D)** Results of Reactome gene set enrichment analysis showing upregulated and downregulated DEGs in the indicated comparison groups.

**(E)** Heatmaps showing the effect of culture condition (2D and 3D) and *MAPK14* genotype on expression of key individual genes involved in cell cycle regulation, growth arrest, senescence and expression of TNF ligands.

### Supplementary Figure 3

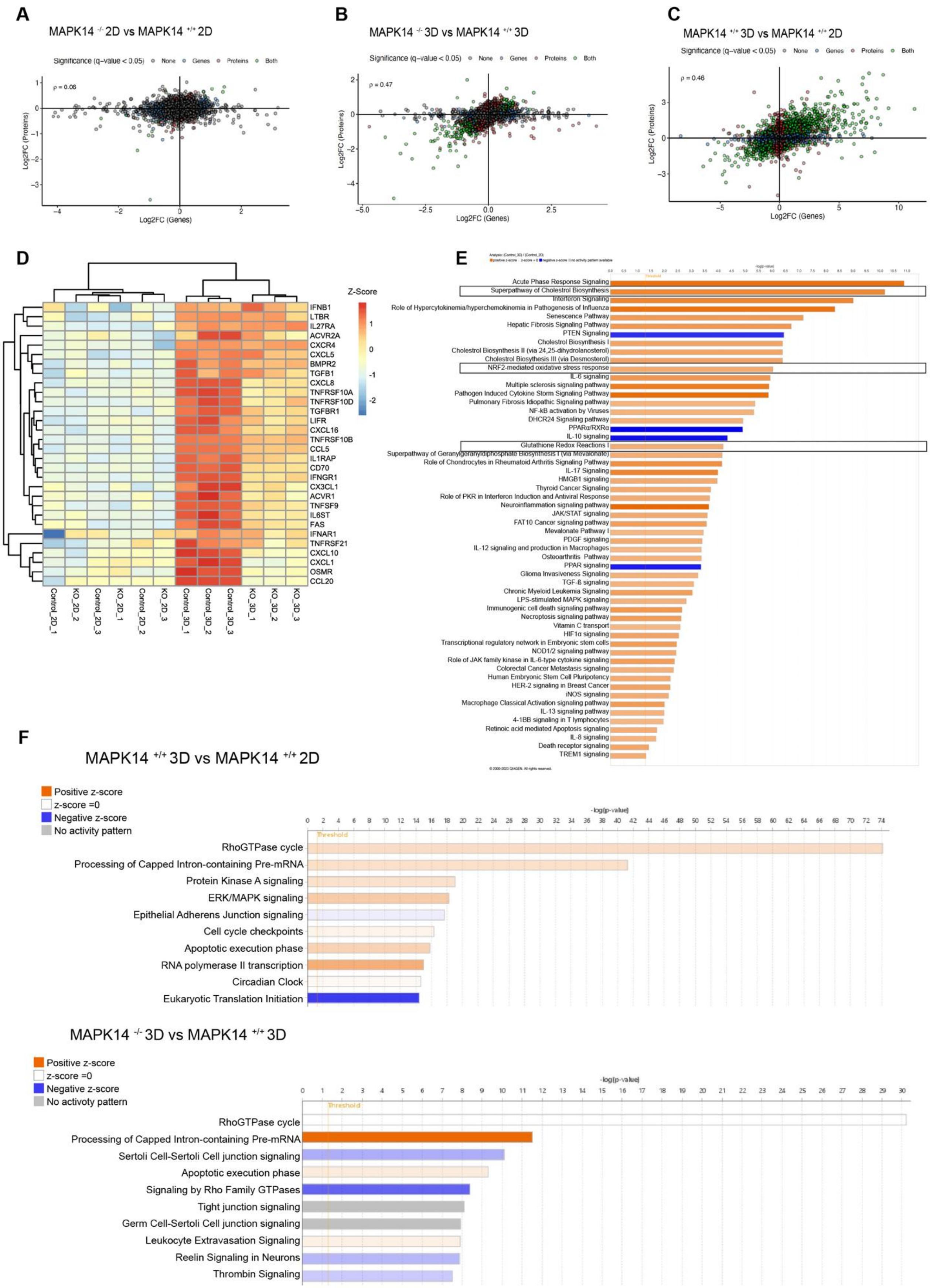

**Supplementary Figure 3. Effect of *MAPK14* on the proteome and phosphoproteome of HEC-50B cells in monolayer and spheroid culture**

**(A-C)** Scatter plots showing the log2 fold change correlation between proteomics and transcriptomics data for the indicated pairwise comparisons of experimental samples. The protein-RNA Spearman correlation coefficient (rho, ρ) values for fold-changes between *MAPK14*$^{-/-}$ 2D vs. *MAPK14*$^{+/+}$ 2D samples, *MAPK14*$^{+/+}$ 3D vs. *MAPK14*$^{-/-}$ 3D samples and *MAPK14*$^{+/+}$3D vs. *MAPK14*$^{+/+}$ 2D samples were ρ= 0.06, ρ= 0.47, and ρ= 0.46 respectively.

**(D)** Heatmap showing differential hierarchical clustering of proteins involved in cytokine and chemokine signaling between *MAPK14*$^{+/+}$ and *MAPK14*$^{-/-}$ HEC-50B cells cultured as monolayers (2D) or spheroids (3D).

**(E)** Bar plots of IPA pathway enrichment of differentially expressed proteins in the HEC-50B 3D cells relative to HEC-50B cells cultured as 2D. Bar plots indicate all significantly enriched (q-score <0.05 and Z score≥1.5) pathways the differentially-expressed proteins may be involved in. Pathways indicated in black boxes highlight differentially regulated metabolic pathways.

**(F)** Bar plots of IPA pathway enrichment in the indicated comparison conditions. Bar plots indicate all significantly enriched (q-score <0.05 and Z score≥2) pathways that may involve the differentially-abundant phosphopetides.

**Supplementary Figure 4**

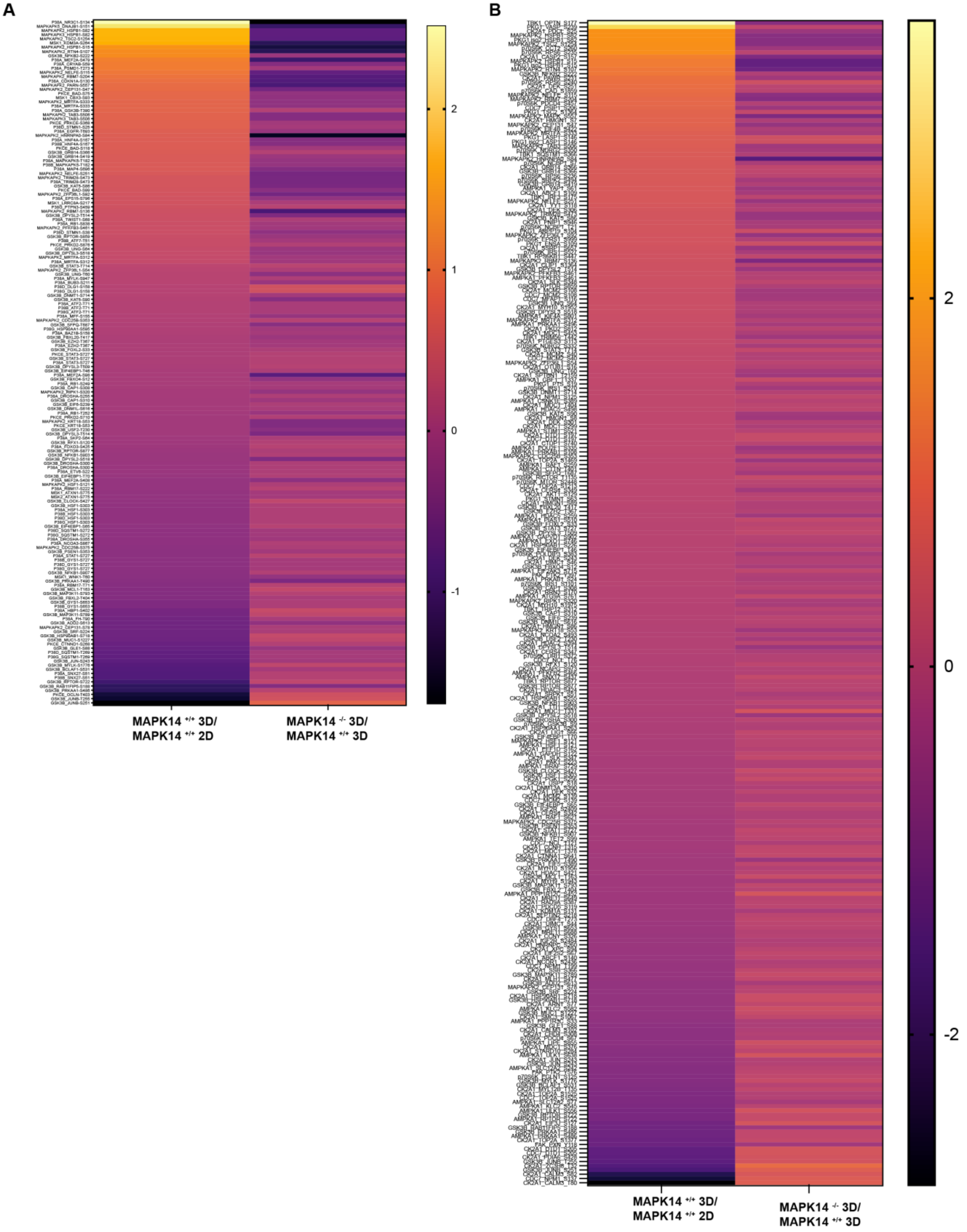

**Supplementary Figure 4. MAPK14 broadly impacts the phosphoproteome of HEC-50B spheroids**

**(A)** Heatmap showing log2FC of a list of known substrates phosphorylated by p38$\alpha$, p38$\beta$, p38$\delta$, and p38$\alpha$-regulated kinases (MAPKAPK2, MAPKAPK3, MAPKAPK5, MSK1, GSK3B and PKCE) as identified using KSEA. The left column indicates fold changes in phosphoprotein substrates that are differentially abundant between *MAPK14*$^{+/+}$ monolayers (2D) and *MAPK14*$^{+/+}$ spheroids (3D). The right column indicates fold changes in substrates that are differentially abundant between *MAPK14*$^{+/+}$ spheroids and *MAPK14*$^{-/-}$ spheroids.

**(B)** Heatmap showing log2FC of a list of substrates for the top 4 and bottom 4 most differentially-regulated protein kinases (identified by KSEA), when comparing *MAPK14*$^{-/-}$ spheroids vs *MAPK14*$^{+/+}$ spheroids. The left column indicates fold changes phosphoproteins that are differentially abundant between *MAPK14*$^{+/+}$ monolayers (2D) and *MAPK14*$^{+/+}$ spheroids (3D). The right column indicates fold changes in substrates that are differentially abundant between *MAPK14*$^{+/+}$ spheroids and *MAPK14*$^{-/-}$ spheroids.

### Supplementary Figure 5

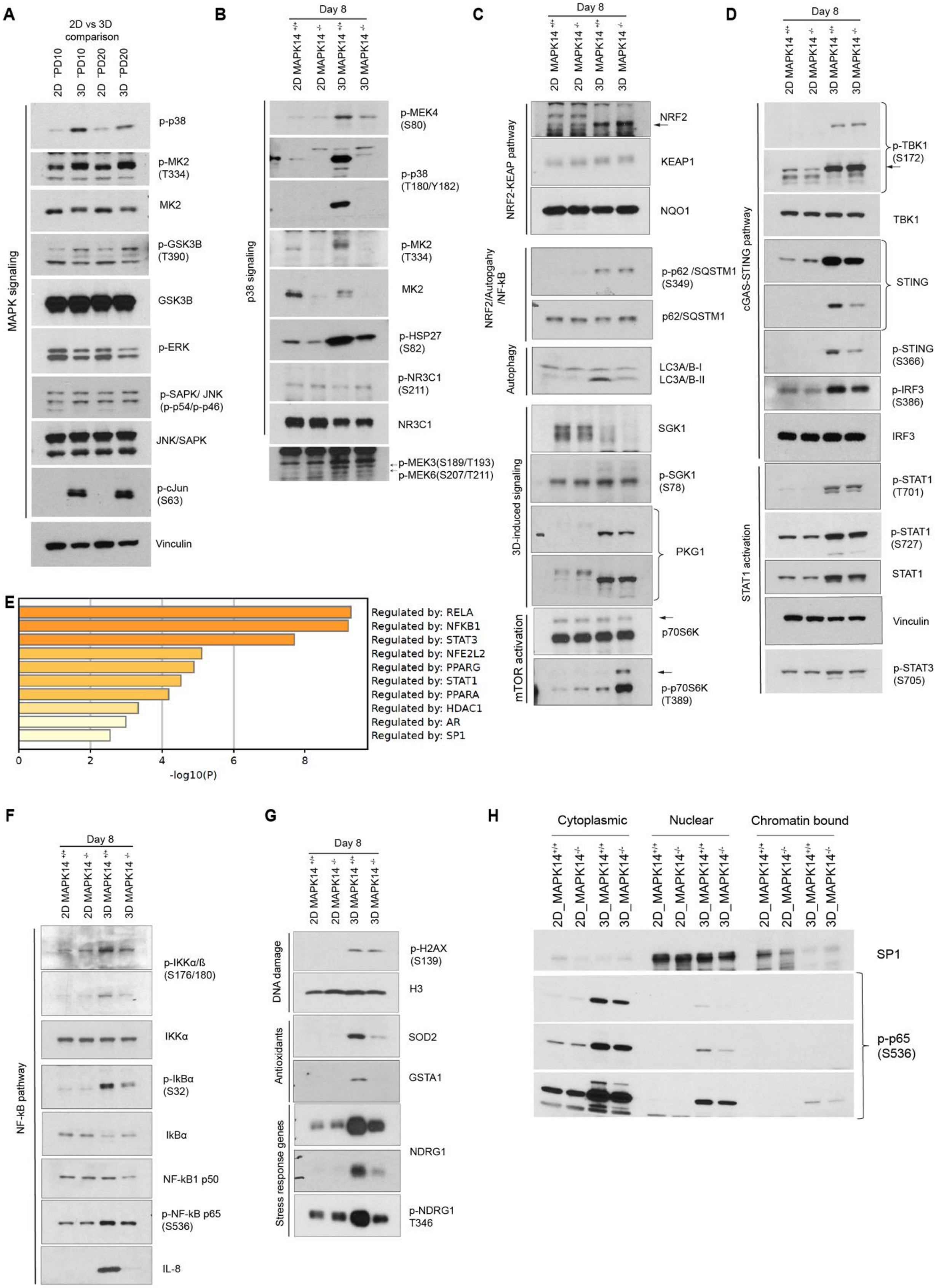

**Supplementary Figure 5. Validation of spheroid culture-induced and *MAPK14*-dependent changes in the phosphoproteome.**

**(A)** Immunoblots showing activation-associated phosphorylation of p38α, its downstream kinases (MK2, GSK3β), and other MAPK family members (ERK, JNK/SAPK) in HEC-50B cells after 10 and 20 population doublings in monolayer and spheroid cultures.

**(B)** Immunoblots showing activation-associated phosphorylation of p38α, MK2, HSP27 in *MAPK14*$^{+/+}$ and *MAPK14*$^{-/-}$ HEC-50B cells after 8 days of growth in monolayer or spheroid culture.

**(C)** Immunoblots showing markers of NRF2, SQSTM1-mediated autophagy, and mTOR pathway activation status in *MAPK14*$^{+/+}$ and *MAPK14*$^{+/+}$ HEC-50B cells after 8 days of growth in monolayer or spheroid culture.

**(D)** Immunoblots showing phosphorylation state and total protein levels of cGAS-STING pathway factors and STAT1 in *MAPK14*$^{+/+}$ and *MAPK14*$^{+/+}$ HEC-50B cells after 8 days of growth in monolayer or spheroid culture.

**(E)** Results of TRRUSTv2 analysis identifying top 10 transcriptional regulators (q-Value <0.05) for genes whose expression (at both mRNAs and protein level) is *MAPK14*-dependent (q-Value <0.05, log2FC$\leq -1$) in HEC-50B spheroids.

**(F)** Immunoblots showing markers of NF-kB pathway activation status and IL-8 levels in *MAPK14*$^{+/+}$ and *MAPK14*$^{+/+}$ HEC-50B cells after 8 days of growth in monolayer or spheroid culture.

**(G)** Immunoblots showing expression of proteins markers associated with of DNA damage (γH2AX), oxidative stress (SOD2, GSTA1) and stress response (NDRG1) in *MAPK14*$^{+/+}$ and *MAPK14*$^{+/+}$ HEC-50B cells after 8 days of growth in monolayer or spheroid culture.

**(H)** Immunoblot showing distribution of p-p65 (S536) between cytoplasmic, nuclear, and chromatin fractions of *MAPK14*$^{+/+}$ and *MAPK14*$^{+/+}$ HEC-50B cells after 8 days of growth in monolayer or spheroid culture.

### Supplementary Figure 6

**A**

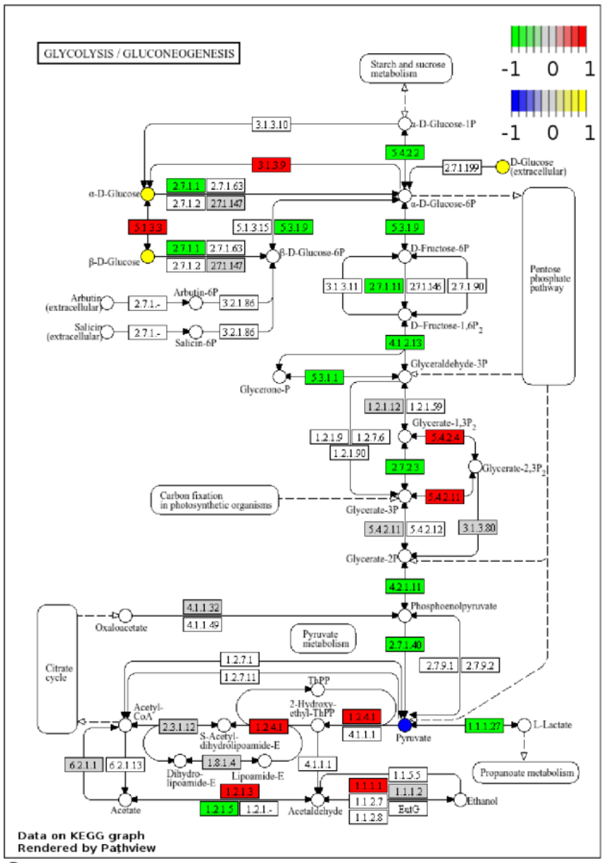


**B**

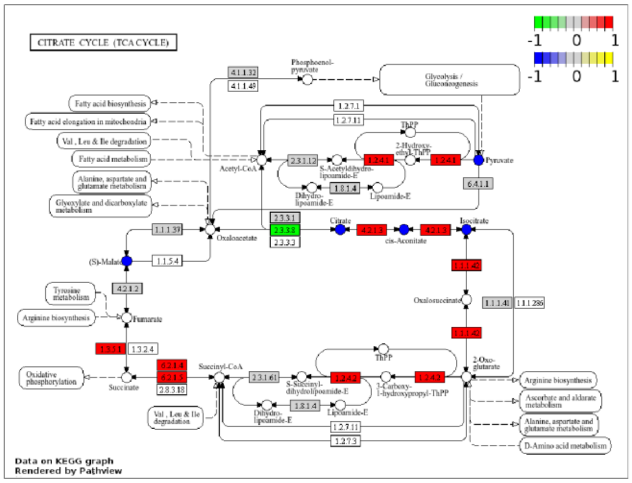


**C**

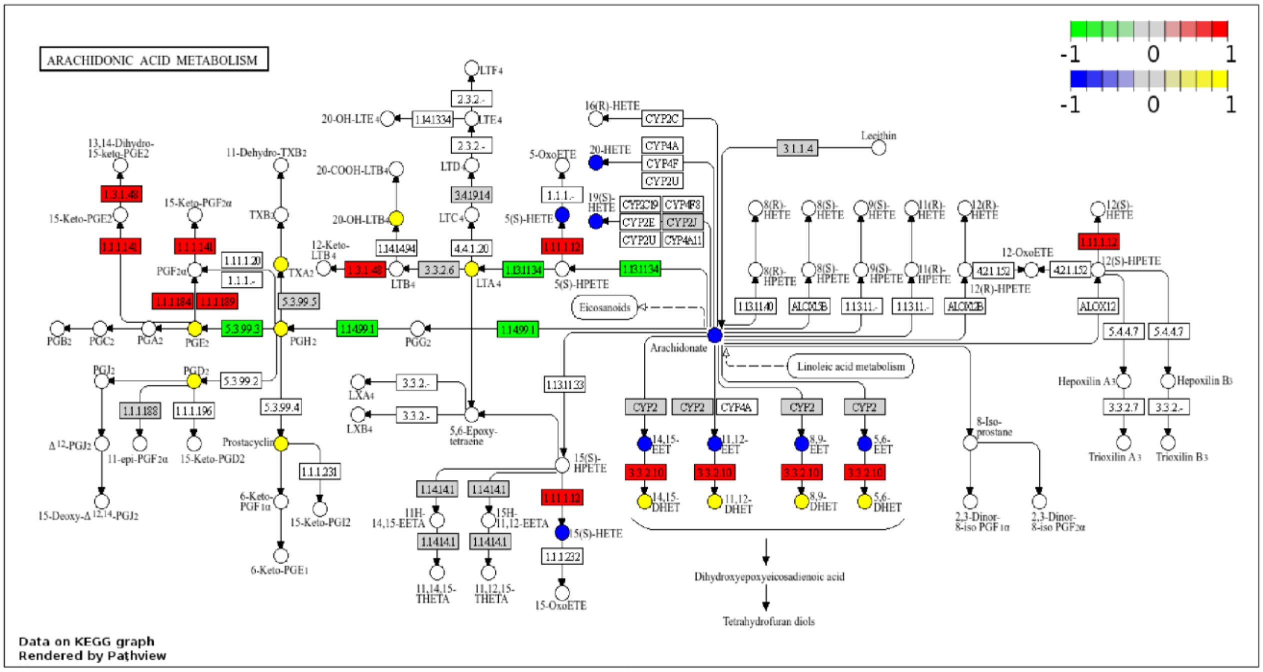

**Supplementary Figure 6. Summary of metabolic pathways impacted by *MAPK14* status.**

Visualization of glycolysis/gluconeogenesis (A), citrate cycle (B), and arachidonic acid metabolism (C) KEGG pathways using metabolites and proteins with $p < 0.1$ between *MAPK14*$^{+/+}$ and *MAPK14*$^{-/-}$ spheroids. mRNAs encoding the enzymes in red are increased in *MAPK14*$^{-/-}$ spheroids, whereas mRNAs encoding enzymes in green are decreased in *MAPK14*$^{-/-}$ spheroids when compared with *MAPK14*$^{+/+}$ spheroids. Metabolites in yellow are increased in *MAPK14*$^{-/-}$ spheroids, whereas metabolites in blue are decreased in *MAPK14*$^{-/-}$ spheroids when compared with *MAPK14*$^{+/+}$ spheroids. Diagrams were made using Pathview.

### Supplementary Figure 7

**A**

Cluster 1

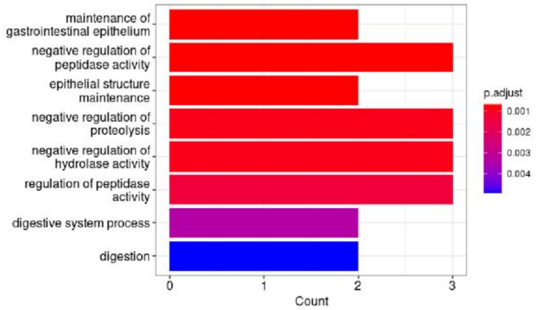


**B**

Cluster 2

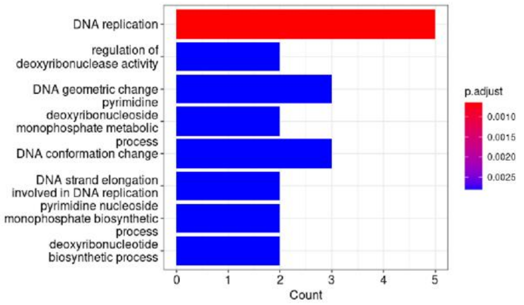


**C**

Cluster 3

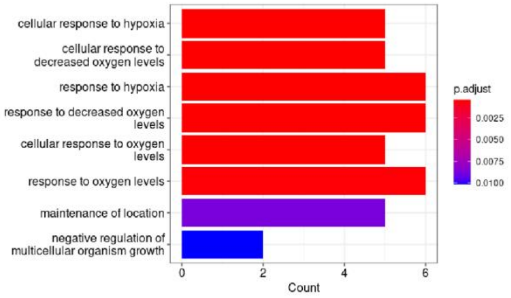


**D**

Cluster 4

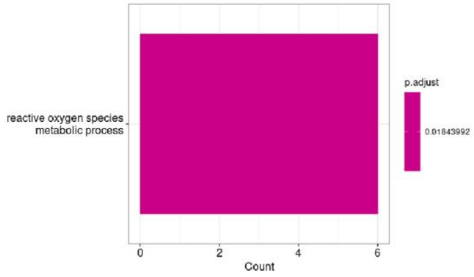


**E**

Cluster 5

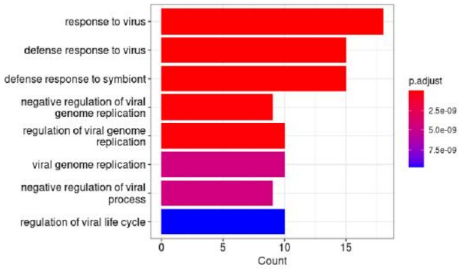


**F**

Cluster 7

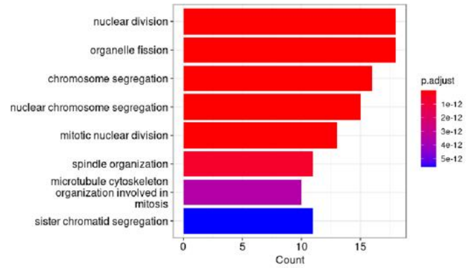


**G**

Cluster 9

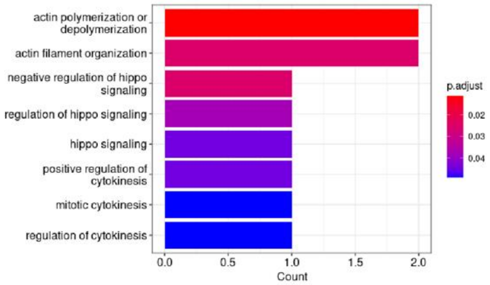


**H**

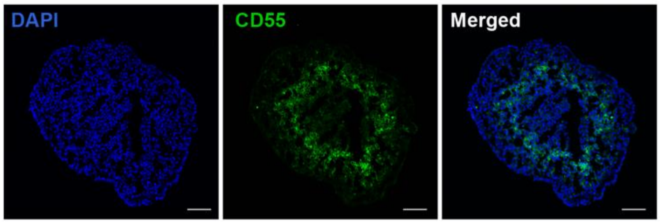

**Supplementary Figure 7. GO analysis of transcriptionally-distinct sub-populations of cells within HEC-50B spheroids**

**(A-G)** GO enrichment analysis for the marker genes expressed by each Seurat cluster from HEC-50B spheroids.

**(H)** Immunofluorescence microscopy of HEC-50B spheroid cryosections showing distribution of CD55-expressing cells at day 8 after seeding. The scale bar represents 100 μm.
